# Rat ultrasonic vocalizations and novelty-induced social and non-social investigation behavior in a seminatural environment

**DOI:** 10.1101/2021.03.16.435680

**Authors:** Indrek Heinla, Xi Chu, Anders Ågmo, Eelke Snoeren

**Author notes:** Corresponding author: Eelke M.S. Snoeren, Current address: Department of Psychology, UiT the Arctic University of Norway, 9037 Tromsø, Norway.

## Abstract

Although rats are known to emit ultrasonic vocalizations (USVs), it remains unclear whether these calls serve an auditory communication purpose. For USVs to be part of communication, the vocal signals will need to be a transfer of information between two or more conspecifics, and with the possibility to induce changes in the behavior of the recipient. Therefore, the aim of our study was to investigate the role of USVs in adult rats’ social and non-social investigation strategies when introduced into a large novel environment with unfamiliar conspecifics. We quantified a wide range of social and non-social behaviors in the seminatural environment, which could be affected by subtle signals, including USVs. We found that during the first hour in the seminatural environment the ability to vocalize did not affect how quickly adult rats met each other, their overall social investigation behavior, their passive social behavior nor their aggressive behavior. Furthermore, the non-social exploratory behaviors and behaviors reflecting anxiety/stress-like states were also unaffected. These results demonstrated that a disability to vocalize did not result in significant disadvantages (or changes) compared to intact conspecifics regarding social and non-social behaviors. This suggests that other (multi)sensory cues are more relevant in social interactions than USVs.

## Introduction

Many animals communicate through vocalization, and the understanding of how and why animals communicate has long been fascinating to scientists [1]. Information encoded by vocal cues has diverse behavioral significance depending on the species. They can, for instance, serve a role in mating rituals, act as warning calls, convey location of food sources, or play a role in influencing the behavior of an interacting partner (reviewed in [2]). The fact that rats can produce vocal signals as audible squeals in the range of 2-4 kHz and ultrasonic vocalizations (USVs, up to ~80 kHz) has been known for a long time [3]. However, researchers are still attempting to understand the structure and function of these calls.

Adult rats emit two main types of ultrasonic vocalizations: the low 22 kHz and the high 50 kHz calls. The 22 kHz calls are assumed to function as alarm calls, since they have been observed mostly in aversive situations/contexts (reviewed in [4]). The 50 kHz calls (ranging between 30-80 kHz), on the other hand, reflecting appetitive calls, are emitted in the presence of for instance a sexual partner and during copulation [5–7], during social [8] and play behavior [9, 10], or after administration of hedonic drugs [11, 12].

Although USVs are reported to be emitted before, during, and/or after certain events, the exact role of these vocalizations to the relevant event is not self-explanatory. Many researchers have proposed that the USVs serve a communicative role, but in order for the vocalizations to be part of *communication*, the vocal signals will need to be a transfer of information between two or more conspecifics, and with the possibility to induce changes in the behavior of the recipient. So far, the empirical evidence remains unsatisfying on whether USVs play a communicative role. Some evidence pointing in the direction of a communicative function is shown by the fact that playback of pre-recorded 22 kHz resulted in defensive behavior of the recipient [13]. With regards to 50 kHz calls, though, different studies have found opposite results. Some have shown that the playback of 50 kHz calls induces transient approach behavior in rats, especially juveniles [14–17], while others have demonstrated that the playback of vocalizations from a conspecific of the opposite sex does *not* induce approach behavior in male nor female adult rats [18, 19]. In addition, it was found that when the emission or receiving of the USVs is disrupted (e.g. by devocalization or deafening), rats hardly elicit different patterns of behavior in their partners [20–23]. Only in juvenile rats, different patterns of play behavior have been found in dyads of silent versus vocalizing rats [24].

In addition, rodents constantly interact with their conspecifics, using different means of communication. Their social behavior consists of more different elements of behaviors besides the approach or play behaviors mentioned above. In a broad sense, social behaviors can be defined as any modality of communication and/or interaction between conspecifics of a given species (see review [25]). Social behavior displayed at the inappropriate time or place or of unfitting intensity can have negative consequences to the individuals even to a social group as a whole. These interactions involve active detection and response to cues from multiple sensory modalities, and a continuous exchange of social information perceived from sensory cues produces an important feedback loop that could change the behavioral responses again. Since the complexity of interactions depends on the potential communication space between individuals, social behaviors are among the most complex behaviors. Unlike some other communication modalities, USV communication has strong directivity, low energy consumption, thus they can be effective over a wide range of distances [26], which makes USVs an interesting candidate for a communicative function in social behavior in rats.

Surprisingly, studies on the role of USVs in social interaction in rats are rare, and the studies that are performed (mainly studying play behavior in juveniles) make use of traditional test settings in which rats are placed in a small arena without the opportunity to express their full repertoire of behavior or interact with multiple conspecifics [24, 27, 28]. As it has been suggested that USVs are used as social-locational cues (providing information about the other conspecifics nearby and their whereabouts) [29], one could argue that for USVs to play a communicative role in social behavior, more space might need to be required than is available in traditional set-ups, for some of these cues to have any significance.

Though, previously we have reported that silencing rats with devocalization procedures did not significantly affect sexual behavior or social interactions, via sniffing behavior, in rat tested in a seminatural environment [21]. As sexual behavior is probably one of the most relevant behavior in which social-locational cues should play a major role, this suggests that USVs do not play an essential role in social interaction. However, the rats in this study were already present the environment for 7 days and were therefore already familiar with each other at the moment of testing. It is hypothetically possible that the rats had already adapted to the communication limitations and modified their interaction behaviors. In addition, individuals with disabled social and communication abilities could perform normally in some situations, whereas, when posed with novel situations, they might experience higher levels of stress and need longer time to adjust to the circumstances. In combination with the idea that appropriate communication and social interaction is probably most important upon first encounter, it would be interesting to look at the role of USVs when rats are introduced to a novel seminatural environment with unfamiliar conspecifics.

Therefore, the aim of our study was to investigate the role of USVs in rats’ social and non-social investigation strategies when introduced into a novel large environment with unfamiliar conspecifics. We quantified a wide range of social and non-social behaviors in the seminatural environment, which could be affected by subtle signals, including USVs. As tracking of the individual’s USVs within a group of rats comes with its own challenges, especially in a large arena, our current study used devocalized and sham-operated vocalizing male and female rats. Another advantage of this approach is that we were able to investigate a batch in which some rats were completely silent. If the emission of USVs plays a role in social investigation behavior, our test conditions should be ideal to detect differences in behavior. Based on our previous findings, we expected that devocalized rats would overall show similar social investigation patterns as sham-operated vocalizing controls in our naturalistic set-up. However, at the same time, we expected that *if* USVs are indeed used as means of communication, it would be most visible during the first encounters with unfamiliar rats. Devocalized rats should then for instance be approached less by others than vocalizing rats.

## Methods

The data was collected from video recordings obtained in a previously performed experiment, resulting in the same materials and methods described previously [21]. The differences between the current and previous study are the behavioral scoring scheme that were used and timing of the observations. In the previous study, the role of USVs in sexual behavior were investigated, while the current study focuses on the role of USVs in other social and non-social behaviors. In addition, in the current study we analyzed the behavior during the first hour after introduction into the seminatural environment when the environment and conspecifics are still novel, whereas the previous study investigated the behaviors on day 7, after they had been familiarized to the new environment.

### Animals

A total of 16 female and 12 male Wistar rats (250–300g upon arrival at an age of ca. 3 months) were obtained from Charles River (Sulzfeld, Germany). Before testing, the animals were housed in same-sex pairs in Macrolon IV open cages (so all the animals were used to hearing vocalizations in the animal room) with tap water and commercial rat pellets available *ad libitum*. All rats had obtained one sexual experience in a copulation test prior to the experiment [21, 30]. The experiment was conducted in accordance with European Union directive 2010/63/EU and was approved by the National Animal Research Authority (ID 5441). The rats were around an age of 3 months at the start of the experiment.

### Surgeries

The procedures were described previously in [21]. Briefly, all females were ovariectomized upon arrival. Operations were done under isoflurane anesthesia and afterwards rats were checked twice daily for 3 days and treated with 0.05 mg/kg buprenorphine every 12 hours (subcutaneously). After obtaining one session of sexual experience two weeks after ovariectomy, seven females and five males were devocalized 3 weeks before they entered the seminatural environment (DEV). Two-centimeter incision was made on the ventral surface of the neck, sternohyoideus muscles were separated and trachea exposed. Next, recurrent laryngeal nerves were cleared from fascia and bilaterally 3mm section of the nerve was removed. The control rats (CTR) received sham surgery (similar procedure, but the nerve was left intact). All animals recovered well from the surgeries.

### Seminatural environment

The seminatural environment (2.4 x 2.1 x 0.75 meters) setup is previously described and illustrated in [31–34]. It consists of a burrow system and an open field area, which are connected by four 8 x 8 cm openings. The burrow system consists of an interconnected tunnel maze (7.6 cm wide and 8 cm high) with 4 nest boxes (20 x 20 x 20 cm) attached, and is covered with Plexiglas. The open area has 75cm high walls, and contains two partitions (40 x 75 cm) to simulate obstacles in nature. A light blocking wall (made of light blocking cloth) between the burrow and the open field allows the light intensity for both arenas to be controlled separately. The burrow system remained in total darkness for the duration of the experiment, while a day-night cycle was simulated in the open area with a lamp 2.5 m above the center that provided 180 lux from 22.45h to 10.30h and approximately 1 lux from 10.30h to 11.00h (the equivalent of moonlight). The light gradually increased/decreased during 30 minutes between 1 and 180 lux.

The floors of both the open area and on the burrow system were covered with a 2 cm layer of aspen wood chip bedding (Tapvei, Harjumaa, Estonia). In addition, the nest boxes were provided with 6 squares of nesting material each (nonwoven hemp fibres, 5 x 5 cm, 0.5 cm thick, Datesend, Manchester, UK), and the open area was equipped with 3 red polycarbonate shelters (15 x 16.5 x 8.5 cm, Datesend, Manchester, UK) and 12 aspen wooden sticks (2 x 2 x 10 cm, Tapvei, Harjumaa, Estonia). Food was provided in one large pile of approximately 2 kg in the open area close to the water supply. Water was available ad libitum in four water bottles.

Two video cameras (VCC-6592; Sanyo, Tokyo, Japan) equipped with a zoom lens (T6Z5710-CS 5.7–34.2 mm; Computar, San Jose, CA, USA) were mounted on the ceiling 2 meters above the seminatural environment to capture the whole environment: one above the open field and another above the burrow system. Infrared lamps provided light for the video camera centered above the burrow.

### Procedure and design

Shortly before (circa 72 hours) being introduced into the seminatural environment, the sham and devocalized males and females were tested for the presence or absence of vocalizations, respectively. As previously described, the male and female rats (who were sexually receptive at this point) were placed in two adjacent chambers covered with sound-absorbing isolation material of extruded polyethylene foam and separated by a wire mesh. A high-frequency sensible microphone (Metris, Hoofddorp, Netherlands) was placed above each chamber and adjusted so that all sounds from within the chamber were recorded, while sounds from the adjacent chamber were not captured by the microphone. The microphone was connected to a computer with the Sonotrack sound analysis system. All devocalized rats used in this experiment did not emit any USV, while the sham animals did. In addition, on another experimental day we have also confirmed the presence of USVs in the seminatural environment.

The day before introduction to the seminatural environment, the subjects’ backs were shaved and tails marked for individual recognition. A rectangle, about 2 x 3 cm, was carefully shaved on the back of the rats the day before introduction into the environment. One female had the rectangle close to the tail, another in the middle of the back, and a third had it close to the neck. The fourth female was not marked on the back. In addition, the tail was marked with one, two, or three transversal, thick black lines. The fourth female was not marked. Males were marked exactly as the females, except that the tail marks were made larger to distinguish between males and females.

Four cohorts of four females and three males were used (resulting in a total number of 9 control females, 7 devocalized females, 7 control males and 5 devocalized males; see Supplementary Tabe 1). Animals in each cohort came from different cages to ensure that they were previously unfamiliar to each other.

Each cohort lived in the seminatural environment for a total of 8 days with full-time recording of all behaviors. After the experiment, the rats were removed from the seminatural environment, the environment was thoroughly cleaned and bedding/nesting materials and food were changed, before a new cohort was introduced.

### Behavioral observation

An experienced observer, blinded for the treatment of rats, scored the behavioral activity of each rat with Noldus Observer XT (Netherlands) during the first 60 minutes after introduction to the seminatural environment. One of 18 different behaviors (see Table 1) was assigned to each rat at any time. Where possible, up to four clarifying modifiers were added: (1) the location where the behavior took place, (2) the partner/recipient of the behavior, (3) if there was a tactile contact with another rat or not and (4) if the given animal initiated the behavior or responded to another rat.

**Table 1.** Description of recorded behaviors.

| Behavior | Description |
| --- | --- |
| Walking/running | Walking or running through the environment |
| Chasing | Running forward in the direction of a conspecific |
| Non-social exploration | Exploring the environment by sniffing, usually when slowly walking or sitting still |
| Interacting with environment | Digging, pushing or carrying bedding/nesting/food material |
| Passive alone | Sitting or sleeping with minimal movement of the head without other rats in close vicinity |
| Passive socially | Sitting or sleeping with minimal movement of the head with at least 1 other rat on maximum 1 rat body length away |
| Hiding alone | Being in the shelter alone |
| Hiding socially | Being in the shelter with at least one other rat |
| Allogrooming | Grooming any part of a conspecific's body, usually on the head or in the neck region |
| Sniffing anogenitally | Sniffing the anogenital region of the conspecific |
| Sniffing nose-to-nose | Sniffing the facial region of the conspecific |
| Sniffing body/head | Sniffing any part of the conspecifics body or head, except for the anogenital and nose region |
| Fighting | Kicking, pouncing, pushing, grabbing, boxing or wrestling another rat |
| Nose-off | Facing another rat and aggressively posturing towards it |
| Self-grooming | Grooming itself |
| Rearing supported | Raising itself upright on its hind paws, facing a wall or an object |
| Rearing unsupported | Raising itself upright on its hind paws, not facing a wall or an object |
| Any other behavior | Behaviors that do not fit any of the other categories (e.g. mounting, drinking, etc) |

### Data preparation and analysis

From the behavioral output file, the following behavioral clusters were generated (see Table 2): *social investigation* (consisting of sniffing anogenitally, sniffing nose-to-nose, sniffing body/head and allogrooming), *non-social investigation* (consisting of walking/running and non-social exploration), *conflict behaviors* (consisting of fighting and nose-off), *passive behaviors* (consisting of passive alone and passive socially), *social passive behaviors* (consisting of passive socially and hiding socially), *non-social passive behaviors* (consisting of passive alone and hiding alone), *all passive behaviors* (consisting of passive alone, passive socially, hiding alone and hiding socially), *hiding* (consisting of hiding alone and hiding socially), *all sniffing* (consisting of sniffing anogenitally, sniffing nose-to-nose and sniffing body/head) and *all rearing* (consisting of rearing supported and rearing unsupported). For these behavioral clusters, but also for the individual behaviors, the latency, frequency, duration and mean duration of episode of each behavior or behavioral cluster for the whole hour in the whole arena were calculated, along with the same parameters separated by location (burrow area versus open area) and into 10-minute time-bins.

**Table 2.** Description of behavioral clusters.

| Cluster | Behaviors within clusters |
| --- | --- |
| Social investigation | Sniffing anogenitally, sniffing nose-to-nose, sniffing body/head, and allogrooming |
| Non-social investigation | Walking/running, non-social exploration |
| Conflict behaviors | Nose-off, fighting |
| Passive behaviors | Passive alone, passive socially |
| Social passive behaviors | Passive socially, hiding socially, |
| Non-social passive behaviors | Passive alone, hiding alone |
| All passive behaviors | Social and non-social passive behaviors |
| Hiding | Hiding alone, hiding socially |
| All sniffing | Sniffing anogenitally, sniffing nose-to-nose, sniffing body/head |
| All rearing | Rearing supported, rearing unsupported |

To further investigate social behavior, it was also calculated how fast each rat met each of their conspecifics, the mean duration of their social interactions, how much time overall did they spend in tactile contact with conspecifics, ratio of social activity (time in social behaviors/overall time), ratio of active non-social behavior (non social investigation/immobility), ratios of different types of sniffing, percentage of unsupported rearing (unsupported rearing/all rearing), and how much time they spent on open arena doing non-social behaviors.

For analysis of the data of the whole hour, a linear mixed model with rat as subject and *treatment* and *sex* as factors was used (IBM SPSS Statistics 26). A modified Benjamini-Hochberg procedure was used (instead of using all possible comparisons, which would yield too strict criteria for behavioral data, we only used p-values of four predetermined clusters: all sniffing, non-social investigation, self-grooming and conflict behavior, in addition to all behaviors with p<0.05) to correct for multiple comparison analysis. The data separated into 10-minute time-bins was analyzed with a repeated measures ANOVA with *time* as a within-subject factor and *treatment* and *sex* as between-subject factors. If Mauchly’s test of sphericity yielded p<0.05, Greenhouse-Geisser test of within-subjects effects is reported, otherwise if Mauchly’s test of sphericity yielded n.s., sphericity assumed test of within-subjects effects is reported.

One devocalized female rat was excluded from the analysis because she spent an overwhelming majority of the time passively (87% of the overall time, in comparison to others with on average 2.8± .3%). The reason remains unclear, but therefore the data throughout the manuscript is presented without this rat.

### Statement Open Science Framework (OSF)

The design of our study was preregistered on OSF on the 17^th^ of December 2019 (https://osf.io/gzkjw). We refrained from the analysis of entry and re-entry latencies of different parts of the environment, because first rats were entered into the environment before starting the videos and therefore we were not able to collect complete data; otherwise there were no changes in analysis.

## Results

Since our data analysis generated a lot of data, only the most relevant findings from the total environment in this section are reported. For more details on different aspects of the data, or the data from the open area and burrow alone, please turn to the supplementary Tables 2-5.

### Social investigation

As mentioned in the introduction, social behavior is a complex behavior that involves multiple aspects. Besides the different categories of social behavior, it also involves the interaction between two or more animals and thus the differentiation in whether a rat is the initiator or responder to a social interaction. To investigate the role of USVs in social behavior, parameters linked to social behavior were explored. The time it took to meet all new conspecifics, the frequency and duration of social behaviors in total as well as initiator or responder, the length of social interaction bouts and the frequency, duration and average time they were being socially investigated were calculated. In addition, it was analyzed how much of these episodes contained actual tactile contact. Interestingly, no differences were found in any of these parameters between silent (DEV) and vocalizing (CTR) female and/or male rats.

First of all, it was found that cohort members met very quickly, as most animals had actively sniffed more than half of their new cohort members within the first minute and had mostly approached all six of their new cohabitants within the first 5 minutes. No differences were found between CTR and DEV animals in terms of latency to approach new conspecifics or being approached by conspecifics (effect of treatment F_(1,23)_=.196; n.s. Fig 1A). In addition, no differences were found between time spent on social investigation behavior (effect of treatment on social investigation F_(1,23)_=.039; n.s. Fig 1B) or its separate subcomponents of social behaviors between CTR and DEV rats (Fig S1A, B, C, Supplementary Table 2). Male rats only spent in general more time sniffing the anogenital or body regions than female rats, but no significant treatment*sex interaction effect was found (see Supplementary Tables 2&3). Similar results were found in the time receiving social investigation behaviors (or its subcomponents) from conspecifics (without necessarily responding to it: effect of treatment on social investigation behavior F_(1,23)_=.007; n.s. Fig 1C) or with regard to the length of the social interaction bouts (effect of treatment: dyads with DEV rat F_(1,23)_<0.01, n.s.; dyads with CTR rat F_(1,23)_=0.81, n.s., DEV-DEV vs CTR-CTR dyads F_(1,23)_=1.923; n.s. Fig S1L, M). Not even when only the first 10 encounters were analyzed separately effect of treatment F_(1,23)_=2.298; n.s. Fig S1H, Supplementary Table 2).

**Figure 1.**
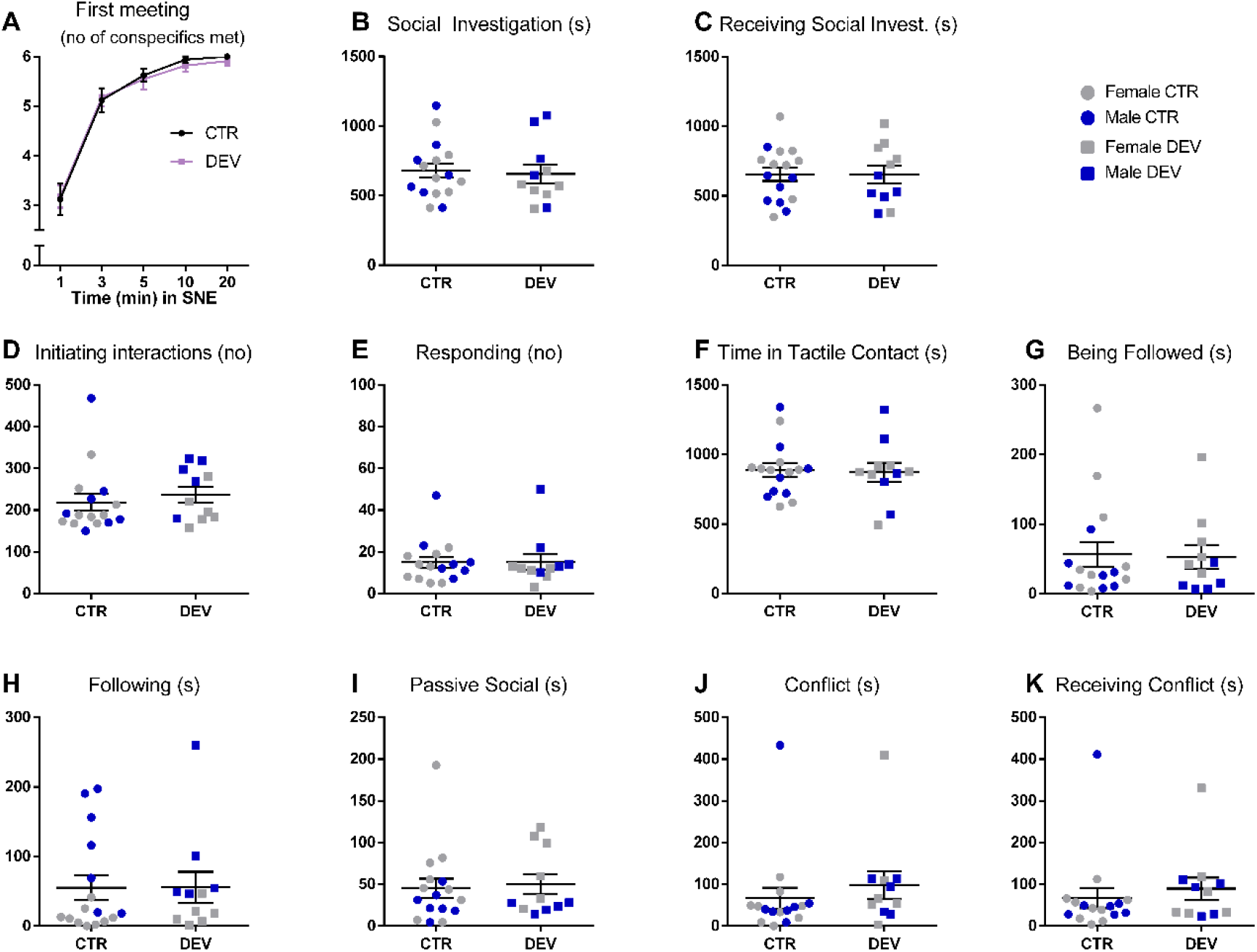
Social behavior of devocalized (DEV, n=11) and sham-operated control (CTR, n=16) rats. (A) The number of different conspecifics that were met within 1, 3, 5, 10 and 20 minutes. (B) Time spent on social investigation (sniffing anogenitally, sniffing nose-to-nose, sniffing body/head, and allogrooming). (C) Time being socially investigated by conspecifics. (D) Number of initiated social interactions. (E) Number of responses to a social interaction initiated by a conspecific. (F) Time spent in tactile contact with other rats. (G) Time being followed by a conspecific. (H) Time spent on following other rats. (I) Time spent on being passive socially (hiding and passive socially). (J) Time spent on conflict behavior (fighting or nose-off) with conspecifics. (K) Time receiving conflict behaviors from conspecifics. Data are shown with individual data points (females in grey, males in blue) with the lines representing the group means. Error bars are representing standard error of the mean SEM. s = seconds, no = number of episodes.

Also when the social behaviors were divided in the episodes in which a rat was the initiator versus the responder or with/without tactile contact, DEV rats initiated (effect of treatment F_(1,23)_=.496; n.s. Fig 1D) and spent a similar amount of time on initiated social behaviors (effect of treatment F_(1,23)_=.218; n.s. Fig S1D) as CTR rats. Similarly, there were no differences in episodes of responding to others (effect of treatment F_(1,23)_=.011; n.s. Fig 1E) or duration of responding to others (F_(1,23)_==.001; n.s.) in social investigation behavior. It should be mentioned, though, that it is sometimes unclear in a seminatural environment which animal initiates the interaction. This limitation was solved by scoring both participants of the social interaction as initiators. Moreover, it was found that the overall time spent *with* tactile contact (effect of treatment F_(1,23)_=,016; n.s. Fig 1F) and the average length of these interactions were not different in CTR and DEV rats (Fig S1I, Supplementary Table 2).

Furthermore, the data revealed no differences in any other behavior involving a conspecific that could have been affected by devocalization, such as following, passive socially and conflict behavior. There was no difference between vocalizing and silent animals in how much time they were being followed (effect of treatment F_(1,23)_=.024; n.s. Fig 1G) or how much they followed others (effect of treatment F=.005; n.s. Fig 1H). Also, when the data at whom they follow (behaviors following DEV and following CTR rats are corrected according to the number of available partners in a given cohort; Fig S1J, K) was analyzed, no significant differences were found. The data analysis of following behavior only revealed a significant sex effects in that female rats were more often being followed (effect of sex F_(1,23)_=4.96; p=.04) and males doing most of the following (effect of sex F_(1,23)_=17.32; p<.001). However, there was no significant interaction effect between treatment and sex (Supplementary Tables 2&3). Additionally, it was found that silent DEV rats spent a comparable amount of time on passive social behavior (and its subcomponents) to vocalizing CTR rats (effect of treatment F_(1,23)_=.085; n.s. Fig 1I), neither were there differences on the time spent on conflict behavior of DEV and CTR rats, neither as an active partner nor as receiving the conflict (effect of treatment as aggressive party F_(1,23)_=.413; n.s. Fig 1J and effect of treatment as recipient F_(1,23)_=.024; n.s. Fig 1K, refer to the Supplementary Table 2 for mean values).

### Non-social investigation and other behaviors

Besides social behaviors, USVs could also affect emotional state of the vocalizing animal itself, which could then influence their non-social investigation patterns in a novel environment or their stress-coping behavior. For example, if USVs had a comforting effect on the rat itself, one could hypothesize that CTR rats might feel safer to explore the novel environment than a DEV rats. Therefore, our study also investigated the non-social investigation strategies of the rats, in addition to parameters like self-grooming, rearing, and time spent in the open area.

However, analysis of the overall time spent investigating the environment (effect of treatment F_(1,23)_=.612; n.s. Fig 2A), in addition to the separate subcomponents walking/running (effect of treatment F_(1,23)_=.30; n.s.) and non-social exploration (effect of treatment F_(1,23)_=.33; n.s.), did not reveal any differences between CTR and DEV rats. There was, though, a sex effect showing that females spent more time on non-social investigation than males (effect of sex F_(1,23)_=14.27; p=.001), but no interaction effect between sex and treatment was found.

**Figure 2.**
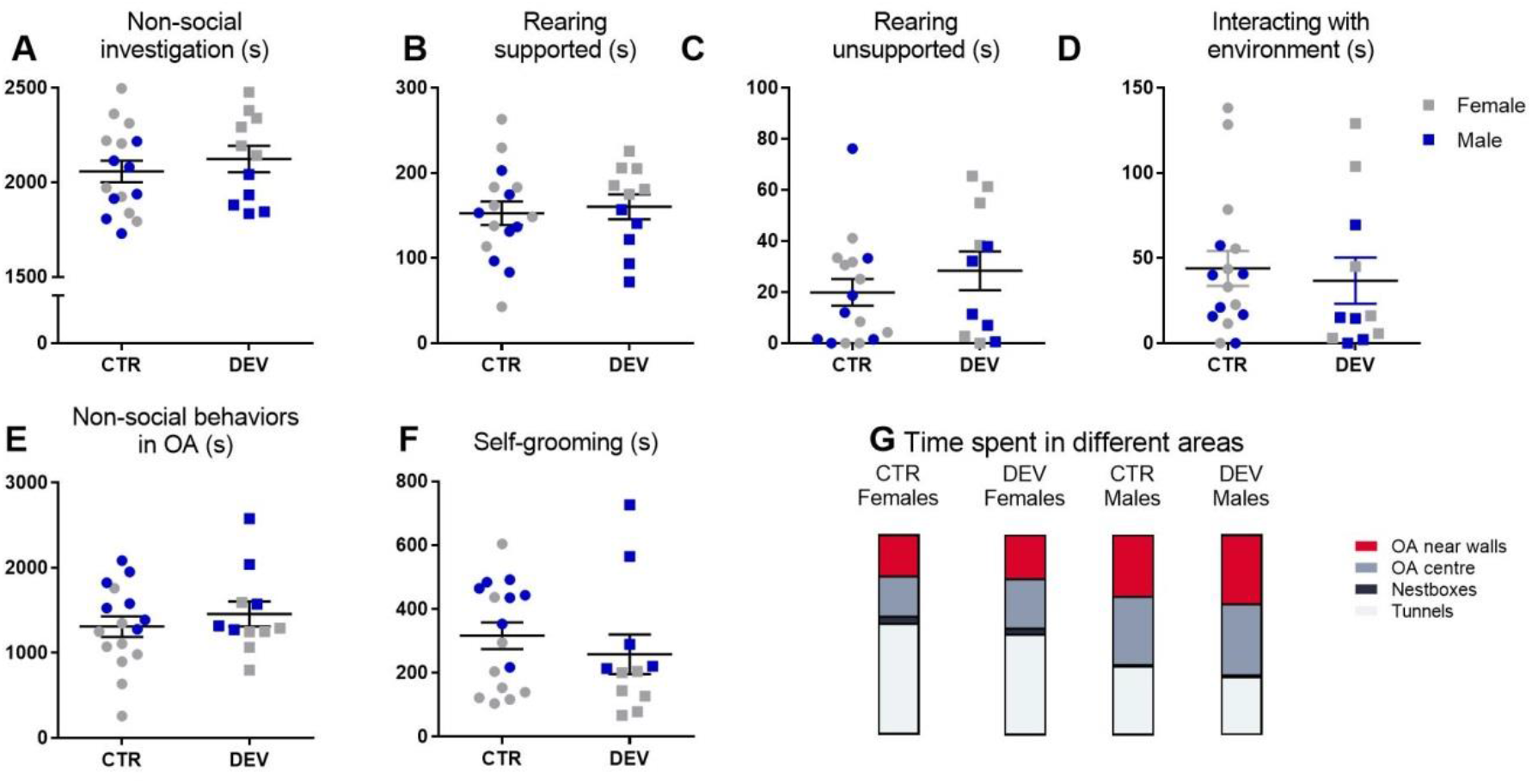
Non-social behavior of devocalized (DEV, n=11) and sham-operated control (CTR, n=16) rats. (A) Time spent on non-social investigation behavior. (B-C) Time spent on rearing suppoted and unsupported. (D)Time spent on interacting with the environment. (E) Time spent in the open area (OA) (excluding social interactions)(F) Time spent on self-grooming. (G) Relative time spent in the different areas of the environment. The height of the colored box represents the proportion of time the rats of the given group on average spent in respective area. Data in A-F are shown with individual data points (females in grey, males in blue) with the lines representing the group means. Error bars are representing standard error of the mean SEM. s = seconds.

Rearing to hind legs provides superior vantage point to investigate the surrounding social and physical environment. A distinction between supported and unsupported rearing was made with the idea that unsupported rearing has been shown to be modulated by anxiety-like states [35]. If emitting USVs would induce a comforting effect, one could assume that CTR rats show more unsupported rearing. In our experiment, however, no effect was found on supported rearing nor unsupported rearing (supported rearing: effect of treatment F=.087; n.s. Fig 2B; unsupported rearing: effect of treatment F_(1,23)_=.703; n.s. Fig 2C). Also when unsupported and supported rearing were combined, no differences between CTR and DEV were found (effect of treatment F_(1,23)_=.267; n.s. Fig S1N), except that females rear more often than males (effect of sex F_(1,23)_=5.786; p=.025).

Other behaviors that could be linked to anxiety-like states and could thus theoretically be affected by USVs if these play a role on emotional state, are behaviors like digging, transporting the bedding material, nesting material and food (combined in the cluster “interaction with the environment”), self-grooming and the time spent in open arena. Data analysis revealed, though, that there were no effects of the absence of USVs on interacting with the environment (effect of treatment F_(1,23)_=.173; n.s. Fig 2D), nor on the amount of time spent in the open arena including (effect of treatmnet F_(1,23)_=.839; n.s.) or excluding the episodes in which they participated in social interactions (effect of treatment F_(1,23)_=.735; n.s.). It was found, though, that male rats spent in general more time in open area compared to females (including social interactions: effect of sex F_(1,23)_=17.008; p<.001; excluding social interaction: effect of sex F_(1,23)_=14.403; p=.001; Fig 2E). Female rats, on the other hand, spent more time in the burrow (tunnels and nestboxes; effect of sex F_(1,23)_=16.432; p<.001), but no effects of treatment were found between CTR and DEV rats (effect of treatment F_(1,23)_=.879; n.s. Fig 2G).

With regard to self-grooming, a potential measure for stress-coping behavior [36–38], male rats self-groomed more (F_(1,23)_=13.68; p=.001) and longer (F_(1,23)_=13.41; p<.001 Fig 2F) than female rats, but no effect of treatment (number of episodes F_(1,23)_=.164; n.s.; time spent F_(1,23)_=.92; n.s.) or interaction effects of sex*treatment were found.

It should be mentioned, though, that anxiety-like states can be accompanied by behavioral inhibition, which can manifest in delayed onset of natural maintenance and exploratory behaviors. But when we compared the latencies to start self-grooming (effect of treatment F_(1,23)_=.337; p=.57, Fig S1O), unsupported rearing (effect of treatment F_(1,23)_=.09;p=.77) or other behaviors (Supplementary Table 4), no differences between CTR and DEV rats were found.

### Behavioral patterns during the course of an hour

At last, it was investigated how the behavioral patterns of the rats changed over the course of the hour to detect if there are any deviations in how devocalized animals habituate to the novel social and non-social environment. Therefore, the data was divided into six 10-minute time-bins and analyzed the behavioral patterns cumulatively.

As expected, some behaviors were performed more or less in the beginning than in the end. The amount of time spent on social investigation (effect of time F_(5,115)_=6.74; p<0.001; Fig 3A), being socially investigated (time effect F_(5,115)_=5.899; p<.001; Fig S2A, Fig S3C&D), and non-social investigation (effect of time F_(3.322,76.399)_=12.03; p<0.001; Fig 3B) slightly decreased over the course of an hour, whereas the time spent on rearing (effect of time F_(5,115)_=2.013; n.s. Fig S2D, especially unsupported rearing: effect of time F_(5,115)_=2.726; p=0.023), self-grooming (effect of time F_(3.039,69.896)_=13.26; p<0.001; Fig S2B), and passive behavior (F_(2.908,66.874)_=4.20; p=0.009; Fig S2C) increased over the course of an hour.

**Figure 3.**
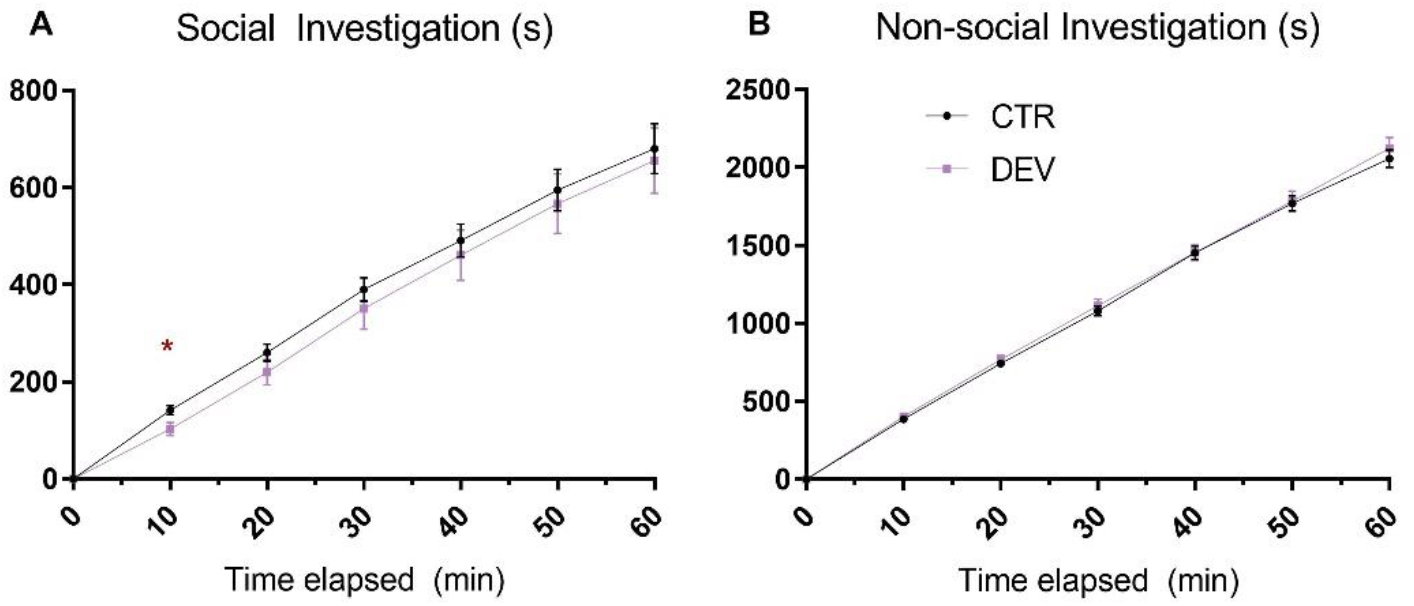
Behavioral patterns during the course of an hour in devocalized (DEV, n=11) and sham-operated control (CTR, n=16) rats. (A) The cumulative time spent on social investigation behavior. (B) The cumulative time spent on non-social investigation behavior. Data are shown in mean±standard error of the mean per 10-miute time-bins. s = seconds, * p<0.05 CTR versus DEV.

Some sex differences were found in these behavioral patterns: male rats showed a steeper decrease in time being socially investigated over the hour (time*sex interaction effect F_(1.223,28.132)_=6.274; p=.014) and a faster increase in self-grooming behavior (effect of time*sex interaction F_(1.419,32.647)_=14.82; p<001) compared to females, while females declined faster in the time spent on non-social investigation (effect of time*sex interaction F_(1.562,35.917)_=13.57; p<001). With regard to rearing, females reared more at the beginning of the experiment and less near the end (effect of time*sex interaction F_(1.621,37.294)_=3.636; p=.045; post-hoc: males vs females for first ten minutes and second ten minutes p=.037; for 40-50 minutes p=.021), while males were initially rearing less with support compared to females (effect of time*sex interaction F_(1.851,42.572)_=5.213; p=.011). But no remarkable interaction effects with treatment (CTR versus DEV) were found.

Only in terms of the amount of time rats spent on social investigation behavior, it was found that DEV rats spent slightly less time on these behaviors within the first 10 minutes compared to CTR rats (p=.012), but this effect disappeared immediately and resulted in an overall lack of interaction effect over the course of an hour (time*treatment interaction F_(1.172,26.949)_=.11; n.s.). Besides, none of the subcomponents of time spent on social investigation showed differences between CTR and DEV rats when analyzed separately. When the data was further divided into 1-minute time-bins, it became clear that the tendency towards a difference in social investigation behavior between CTR and DEV rats occurs in the minutes between 3 and 12 (Fig S3A&B), after which the DEV rats catch up again with the CTR rats.

With regard to rearing, there was no overall time*treatment effect (effect of time* treatment interaction F_(1.621,37.294)_=.03; n.s.). However, silent rats did rear significantly more within the first ten minutes compared to vocalizing rats (p=.007). This effect was probably caused by supported rearing (p=.006). Further analysis into 1-minute time-bins revealed that the difference in supported rearing between CTR and DEV rats was present around the 1^st^ to 10^th^ minute, after which they show comparable amount of rearing again (Fig S3E&F).

A summary of the main findings described below can be found in Table 3.

**Table 3.** Summary of main findings.

|  |  |
| --- | --- |
| No effects were found between CTR and DEV rats on the following parameters of social behaviors: |  |
| - | latency to approach new conspecifics |
| - | time spent on social investigation behavior |
| - | time receiving social investigation behavior from conspecifics |
| - | length of social investigation bouts |
| - | time spent on social behavior as initiator or responder |
| - | overall time spent with tactile contact |
| - | average length of social interactions |
| - | time spent on social passive behavior |
| - | time spent on conflict behavior |
| - | time spent on following behavior |
| No effects were found between CTR and DEV rats on the following parameters of non-social behaviors: |  |
| - | overall time spent investigating the environment |
| - | time spent rearing |
| - | time spent interacting with the environment |
| - | time spent self-grooming |
| - | time spent in open area during non-social behaviors |

## Discussion

In our study, we investigated the role of USVs in social interactions and non-social investigation of a novel environment with unfamiliar conspecifics in adult rats. Our findings show that silent and vocalizing rats behave very similarly in the first hour of exposure to the new environment. We found no differences in social interaction and non-social investigation behaviors between sham and devocalized rats. Silent rats spent comparable amount of the time on social interactions as vocalizing rats, independent of whether they were the initiator or the receiver. In addition, silent and vocalizing rats also familiarized in the same way with new neighbor conspecifics and novel environment, respectively.

This is in line with our hypothesis that was based on the findings of previous studies in which devocalization did not have an effect on sociosexual behavior with familiar rats [20–23]. Interestingly, though, our study is also in line with another recent study by Redecker et al. who have studied the social behavior and USV production of heterozygous (Cacna1c+/-) and wildtype (Cacna1c+/+) rats [27], a genetic modification of calcium voltage-gated channel subunit that have been linked to deficits in social behavior in mice [39]. Upon their expectations, they found that Cacna1c+/- rats emitted less USVs during social interactions than the controls. However, although their auditory cues were reduced, the rats, both mutated and wild type, did not show any differences in social behavior, measured as sniffing, following, social grooming and crawling under/over [27]. Our studies therefore both show that a reduction (or depletion) of USVs does not affect social interaction behavior.

Our experiment revealed that the emission of USVs did not affect rats’ approach behavior in the seminatural environment. This is somewhat contradicting with previous studies that have shown a temporary approach behavior to the playback of 50 kHz calls [14, 15, 40, 41]. However, we have not been able to replicate these findings on approach behavior even in a smaller arena [18, 19], and when rats were able to choose between an intact or devocalized conspecific, the silent rats were just as much approached and preferred as play or sexual partner as the vocalizing rats in traditional test settings [6, 19, 40, 42]. Therefore, it remains unclear what the function and significance of this sort of short-lasting approach behavior is.

*If* USVs indeed modulate rats’ social interactions and induce approach behavior, it would mean that rats that are incapable of vocalizing should be approached less than intact conspecifics. Additionally, *if* USVs could act as a reinforcer of the behavior, the bouts of social interaction between two vocalizing rats should last longer than bouts between dyads from which one or both are devocalized. Another possibility could be that differences would have been found in the approach behavior towards which part of the body (anogenital region, body, nose) is targeted in devocalized and vocalizing rats. Consequently, if USVs played a role in modulating rats’ social interactions, vocalizing rats should perform and/or receive more interactions compared to devocalized rats. However, in our experiment, there were no differences in how quickly devocalized and vocalizing control rats met their cohort members nor in any parameters regarding approach. Even though the vocalizing animals showed a tendency towards increased social investigation early in the experiment, devocalized animals displayed comparable amount of social investigation at the beginning and throughout the hour. It seems that the transient approach behavior, which has been reported in several playback studies, is not visible in a more naturalistic settings with adult rats. Interestingly, these previously reported approach behaviors have only been seen once per each individual and lasted only short, which would then be in line with our findings. Since the effect on approach has previously been found strongly in juvenile rats, and we have indeed replicated this approach (not published), it is also possible that the role of USVs could differ during lifetime. In general, social behavior of the adult rats in our experiment was not affected by their own nor their partner’s ability to vocalize. This supports the idea that USVs do not play a significant role in modulating communication in adult rats.

This conclusion makes us wonder what the function of these calls then could be. Could, for instance, the ability to vocalize modulate rat’s own or partner’s emotional state? One study has shown how rats, that have been trained to react to different sounds to either earn a positive reward (sucrose) or to avoid unpleasant loud white noise, treat an ambiguous cue as positive (predicting reward) if it is preceded by the playback of 50 kHz calls and treat similar cue as negative (predicting unpleasant white noise) if preceded by 22 kHz calls [43]. This implies that 22 kHz and 50 kHz calls are indeed involved in inducing negative and positive responses. In our study, however, we investigated whether potential feedback from vocalizing rat would change dynamics of the social interaction such as length of the interaction, preference for tactile contact or escalation to aggression. Interestingly, in the study by Redecker et al. it was found that the Cacna1c+/- rats, who have reduced USVs emission, did spend more time in physical contact than the Cacna1c+/+ rats [27]. However, our findings did not show any signs of changes in physical contact, type of contact or escalation to aggression upon devocalization. The differences in results could then be explained by the use of a large seminatural environment in which rats are able to choose the type of interaction they prefer in the moment, instead of being forced into a certain behavior.

Another possibility is that vocalizing itself can have a comforting effect on a rat and that devocalization could thus influence their stress-coping and/or non-social investigation behavior in the novel environment. For example, if USVs modulate anxiety/stress-like states of the emitter, one could hypothesize that vocalizing rats should be more comfortable to initially explore the environment, rear more frequently, and spend more time in anxiogenic parts of the enclosure, and/or self-groom less than devocalized rats with less experiencing the comforting effect of emitting USVs. Such self-comforting effect was indeed reported in the study of Cacna1c+/- rats, as the mutated animals self-groom more, show less digging behavior and rearings when interacting in pairs than the control rats [27]. In the current context, however, we again found no differences between vocalizing and silent rats in terms of self-grooming or manipulating the environment (including digging). This does not necessarily contradict the previous findings, since the knockout strain Cacna1c+/− rats with a dysfunctional calcium voltage-gated channel subunit alpha 1 C could have different underlying reasons for this change in behavior. However, our findings at least suggest that the emission of USVs does not modulate stress-related behaviors. At the same time, it is still possible that USV emission is initiated by the same internal state that also facilitates the given behaviors, something we would not be able to see in our devocalized rats. The emitted USVs could then also in theory be a by-product of the given behaviors, which would then indicate that a change in these behaviors result in a reduced number of USVs, but not the other way around.

In our data, we did find an initial increase in supported rearing in devocalized rats. Our exploratory methods are not suitable to explain whether this increase in exploratory rearing is related to the reduced social investigation in the same time window (they can only perform one behavior at the same time), and could then just as well be explained as an unfortunate artifact. Unsupported rearing, which is linked to susceptibility to acute stress [35], was not affected by devocalization. Along with the lack of effects in the other behavioral parameters that could reflect anxiety/stress-like states such as self-grooming (time spent and latency to start) and the time spent in the anxiogenic parts of the environment, our data suggests that the ability to vocalize does not modulate rats’ anxiety/stress-like states.

Previously, we and others have suggested that this could mean that USVs may be purely a by-product of the arousal linked to the behaviors [30, 44, 45], and that 50-kHz USVs are just a by-product of locomotion and breathing. It should be mentioned, though, that the advances in research techniques have now made it possible to study this possibility in more detail and resulted in the conclusion that USVs are not just simply a byproduct. Evidence showed that the emitted USVs are indeed tightly linked to locomotion [29], breathing [46] and cardiovascular function [45], and they are even interlocked with active sniffing [47]. However, the fact that they also actively sniff without the emission of USVs [47], and can both vocalize without movement and move without vocalizing [29] weakens the argument for a by-product effect. Besides, the vocal production apparently increases before locomotion begins [29], and a new call type can be started at any point during the exhalation phase [46, 48, 49]. It should be taken into account, though, that *if* USVs are more than just a by-product of arousal, there should be more information in nuances of the vocal communication, as they have an extensive USV ‘vocabulary’ [50, 51]). So far, many studies have neglected the existence of this vocabulary, and the possible role different type of calls, and the sequence of calls, must have if USVs serve a communicative role after all. Moreover, some studies in juvenile rats that did investigate different types of calls did not find an one-to-one relationship between any movements and specific vocalizations, nor did they show a direct change in the conspecific’s behavior [10, 52]. Thus, altogether we conclude that USVs are unlikely functioning for communication, neither are they involved in regulating non-social exploring behaviors.

It is important to mention, though we failed to find USVs’ effect, that our study does not *exclude* the possibility that vocalizations play a communicative role in social and non-social behavior. It could simply be the case that other (multi)sensory cues are more relevant in these interactions, and compensate for the lack of vocalizations, something that we have shown before with approach behavior in a sexual context [53]. It could still be possible that if rats are never exposed to auditory stimuli, they would fail to socially interact normally. It was shown by Kisko and colleagues that not only devocalized juvenile rats played less, but also that intact rats housed with devocalized rats showed reduced levels of play behavior [24]. They suggested that rats could have a critical period in which the lack of exposure to vocalizations could determine their behavior later in life. This is an interesting theory that should be explored in the future, but our findings at least support the notion that vocalizations are not the most essential way of communication later in life.

In conclusion, our data show that devocalized adult rats do not show altered social interaction behaviors due to their inability to vocalize. Silent and vocalizing rats show similar patterns and types of social interactions, and do not use other social and non-social investigation strategies when introduced to a novel environment with unfamiliar conspecifics. Our data, therefore, does not provide any evidence that USVs play a communicative role in social behavior, nor do they serve a role in regulating non-social investigation behaviors. Although it cannot be excluded that USVs play some unrevealed role in social behavior, it is clear that other non-USV sensory cues are more relevant in these interactions and could have compensated for the lack of vocalizations. New interesting research techniques using complex algorithms to link behaviors to distinct pattens of USVs, as those used nowadays for mice [54], are needed in the future to explore the potential role of USVs in social behavior in naturalistic environments.

## Acknowledgements

We also would like to thank Ragnhild Osnes, Carina Sørensen, Nina Løvhaug, Katrine Harjo, and Remi Osnes for their excellent care of the animals.

