## Supplemental materials for "Rat ultrasonic vocalizations and novelty-induced social and non-social investigation behavior in a seminatural environment"

Corresponding author: Eelke M.S. Snoeren

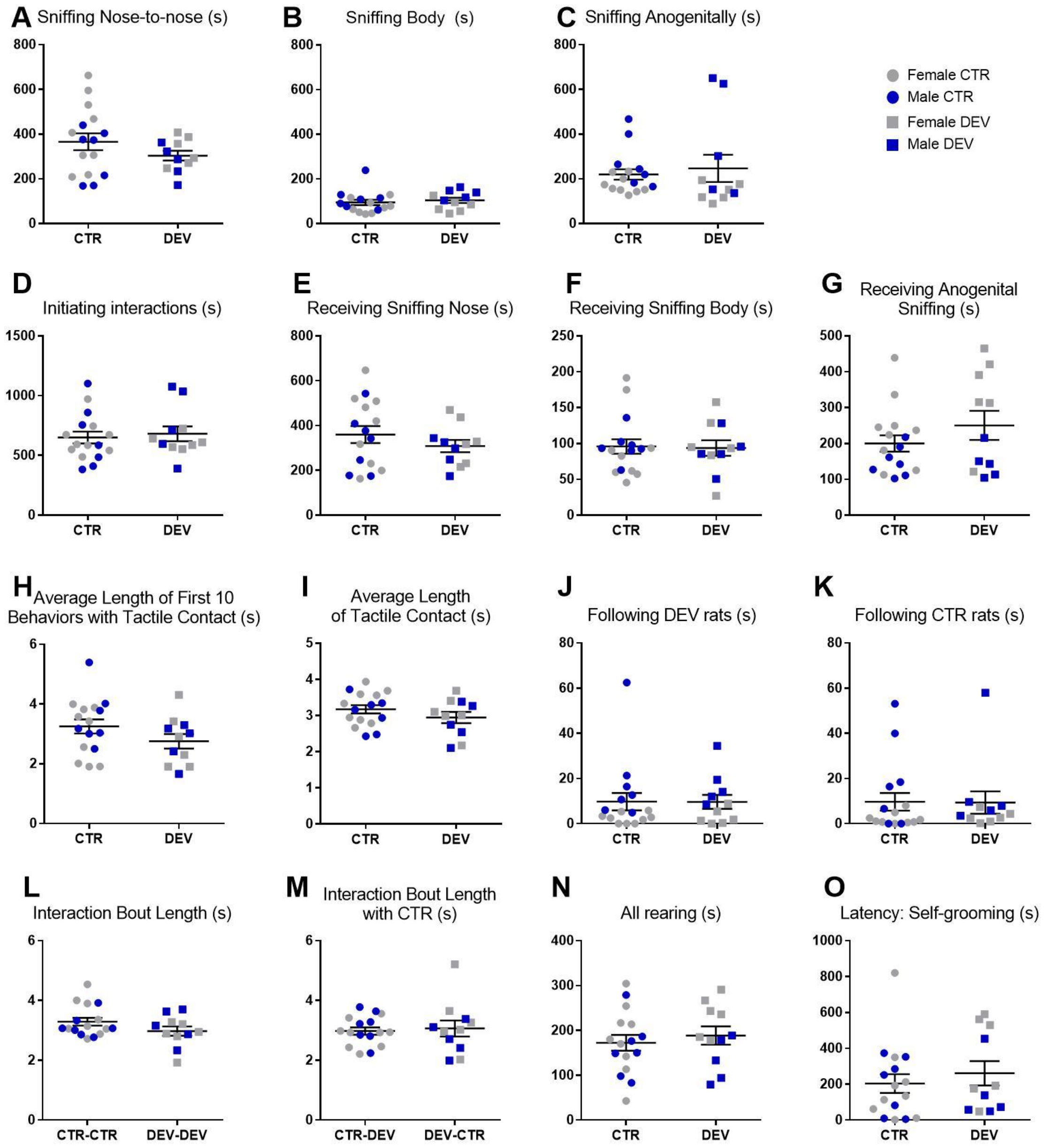


***Figure S1.*** *Social and non-social behavior of devocalized (DEV, n=11) and sham-operated control (CTR, n=16) rats. (A) Time spent sniffing the nose area of a conspecific. (B) Time spent sniffing the body/head area of a conspecific. (C) Time spent sniffing the anogenital region of a conspecific. (D) Time spent on initiated social interactions. (E) Time being sniffed on the nose area. (F) Time being sniffed on the body/head area. (G) Time being anogenitally sniffed. (H) Average length of the first 10 behavioral episodes with tactile contact. (I) Average length of the behavioral episodes with tactile contact. (J) Time spent following devocalized rats. (K) Time spent following vocalizing (sham) rats. (L) Average length of a social interaction episode. (M) Average length of a social interaction episode with a sham rat. (N) Time spent rearing. (O) Latency to start self-grooming. Data are shown with individual data points (females in grey, males in blue) with the lines representing the group means. Error bars are representing standard error of the mean SEM. s = seconds*

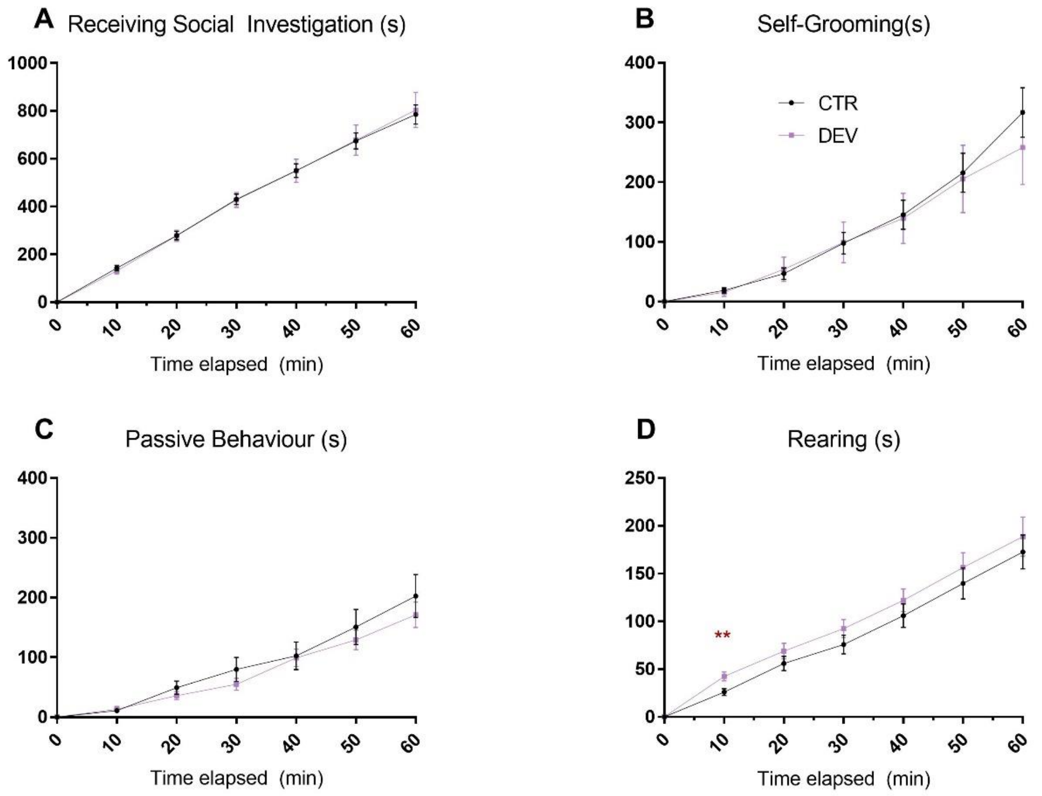


**Figure S2.** *Behavioral patterns during the course of an hour in devocalized (DEV, n=11) and sham-operated control (CTR, n=16) rats. (A) The cumulative time being socially investigated. (B) The cumulative time spent on self-grooming. (C) The cumulative time spent on passive behaviors. (D) The cumulative time spent on rearing. Data are shown in mean±standard error of the mean per 10-miute time-bins. s = seconds, * p<0.05 CTR versus DEV.*

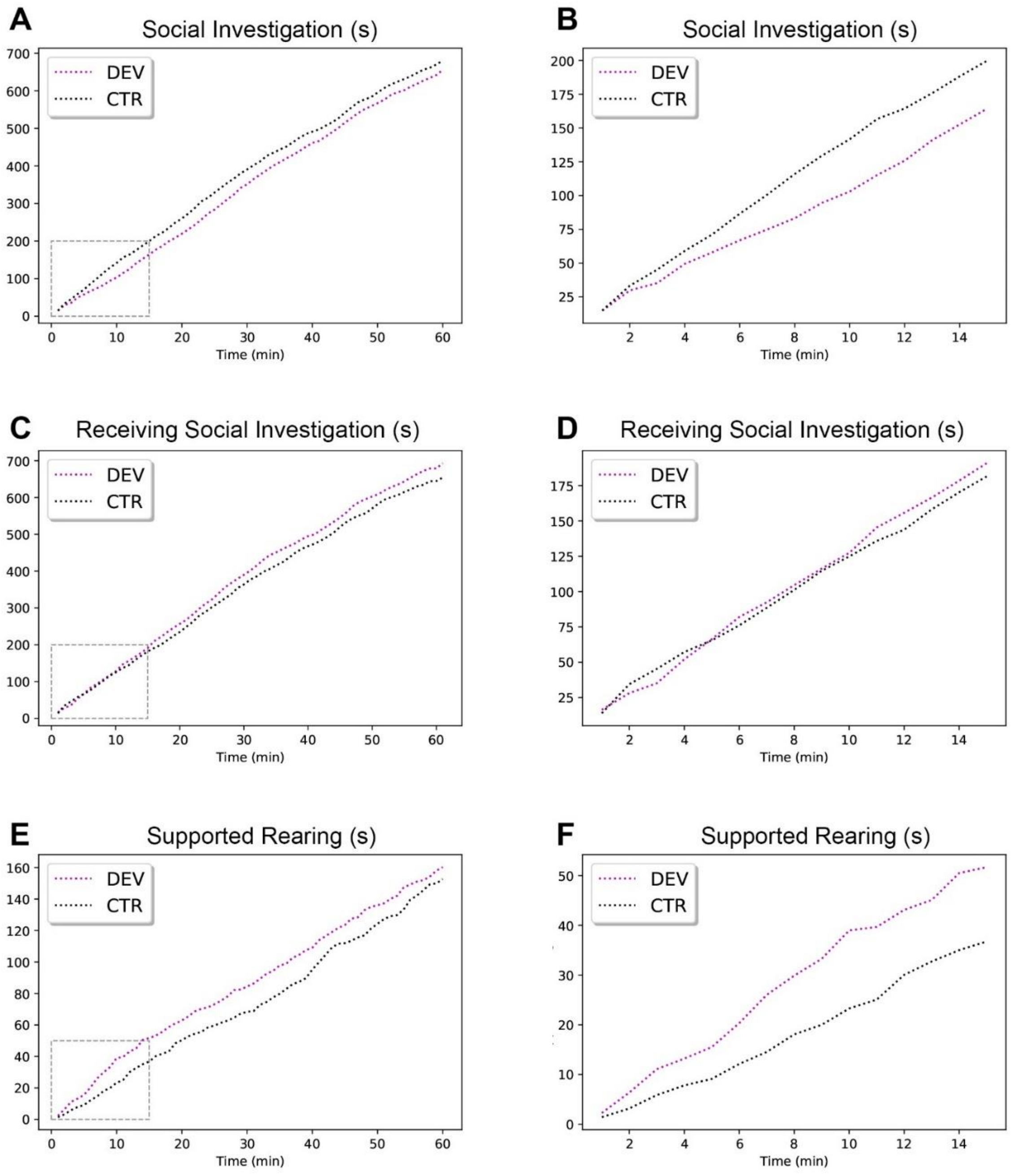


***Figure S3.*** *Behavioral patterns during the course of an hour (A,B,E) or the first 15 minutes (B,D,F) in devocalized (DEV, n=11) and sham-operated control (CTR, n=16) rats. (A,C,E) The cumulative time spent on social investigation behavior, being socially investigated and supported rearing, respectively. (B,D,F) Enlargement of the squares of figure A,C,E respectively, which represents the cumulative time spent on social investigation behavior, being socially investigated and supported rearing for the first 15 minutes. Data are shown in mean over 10-miute time-bins (A,C,E) or 1-minute time-bins (B,D,F). s = seconds*

Supplementary Table 1. Rats in cohorts

| Cohort | Females | Males |
|---|---|---|
| I | F10 – DEV<br>F12 – CTR<br>F14 – CTR<br>F16 – DEV | M10 – CTR<br>M12 – CTR<br>M13 – DEV |
| II | F4 – DEV<br>F19 – CTR<br>F13 – DEV<br>F15 – CTR | M2 – DEV<br>M4 – CTR<br>M6 – DEV |
| IV | F2 – CTR<br>F31 – CTR<br>F6 – CTR<br>F33 – DEV | M8 – CTR<br>M14 – DEV<br>M15 – CTR |
| V | F9 – CTR<br>F11 – CTR<br>F32 – DEV<br>F34 – DEV | M7 – CTR<br>M9 – CTR<br>M11 – DEV |

DEV – Devocalized, CTR – sham operated, vocalising

Cohort III was not included in this study due to problems with video recordings

Supplementary Table 2

Duration — Behavior in the whole environment

| | DEV | CTR | Females DEV | Females CTR | Males DEV | Males CTR | Effect of treatment F | Effect of treatment p* | Effect of sex F | Effect of sex p | Interaction effect F | Interaction effect p |
|---|---|---|---|---|---|---|---|---|---|---|---|---|
| Social investigation | **655.8** ± 64.5 | **680.3** ± 49.9 | 546.9 ± 33.6 | 663.0 ± 57.6 | 786.6 ± 110.6 | 702.4 ± 86.2 | 0.04 | 0.85 | 2.95 | 0.09 | 1.52 | 0.23 |
| Non-social invest | **2123.3** ± 66.6 | **2057.8** ± 55.1 | 2304.2 ± 45.7 | 2124.9 ± 78.7 | 1906.3 ± 34.1 | 1971.4 ± 61.0 | 0.61 | 0.44 | 14.27 | 0.00 | 2.80 | 0.11 |
| Immobile+hiding | **171.0** ± 20.4 | **202.5** ± 34.8 | 195.9 ± 20.3 | 274.7 ± 48.4 | 141.2 ± 33.0 | 109.7 ± 16.6 | 0.34 | 0.57 | 7.31 | 0.01 | 1.84 | 0.19 |
| All Rearing | **188.6** ± 19.5 | **172.6** ± 17.1 | 233.5 ± 16.7 | 182.1 ± 24.4 | 134.8 ± 19.6 | 160.3 ± 22.6 | 0.27 | 0.61 | 5.79 | 0.03 | 2.36 | 0.14 |
| Conflict | **98.1** ± 31.6 | **66.3** ± 24.7 | 115.9 ± 55.0 | 45.2 ± 11.5 | 76.8 ± 17.1 | 93.4 ± 52.6 | 0.41 | 0.53 | 0.01 | 0.91 | 1.08 | 0.31 |
| All Sniffing | **654.6** ± 64.5 | **680.2** ± 50.0 | 546.9 ± 33.6 | 663.0 ± 57.6 | 783.9 ± 111.2 | 702.2 ± 86.2 | 0.05 | 0.84 | 2.88 | 0.10 | 1.48 | 0.24 |
| All Passive | **82.1** ± 6.1 | **114.4** ± 19.1 | 77.1 ± 5.4 | 141.5 ± 27.9 | 88.2 ± 11.3 | 79.5 ± 17.6 | 1.40 | 0.25 | 1.17 | 0.29 | 2.40 | 0.14 |
| All hiding | **88.9** ± 19.5 | **88.2** ± 29.9 | 118.7 ± 19.2 | 133.3 ± 47.4 | 53.0 ± 28.8 | 30.1 ± 9.7 | 0.01 | 0.92 | 4.79 | 0.04 | 0.24 | 0.63 |
| Non-soc passive | **120.9** ± 17.9 | **157.2** ± 27.5 | 122.8 ± 16.2 | 214.9 ± 37.3 | 118.6 ± 34.1 | 82.9 ± 15.5 | 0.72 | 0.40 | 4.22 | 0.05 | 3.72 | 0.07 |
| Social Passive | **50.1** ± 11.3 | **45.3** ± 11.1 | 73.1 ± 15.3 | 59.8 ± 17.8 | 22.5 ± 2.3 | 26.7 ± 5.5 | 0.09 | 0.77 | 7.29 | 0.01 | 0.32 | 0.58 |
| Walking/running | **1906.7** ± 63.3 | **1857.3** ± 54.5 | 2049.9 ± 59.9 | 1912.7 ± 78.5 | 1734.9 ± 58.2 | 1786.0 ± 63.7 | 0.30 | 0.59 | 7.93 | 0.01 | 1.44 | 0.24 |
| Interacting with the environment | **36.7** ± 12.9 | **44.0** ± 9.9 | 50.5 ± 20.1 | 56.8 ± 15.5 | 20.2 ± 11.3 | 27.4 ± 6.8 | 0.17 | 0.68 | 3.35 | 0.08 | 0.00 | 0.98 |
| Following | **55.9** ± 21.1 | **55.2** ± 16.9 | 17.4 ± 6.1 | 12.9 ± 4.1 | 102.1 ± 36.4 | 109.6 ± 26.6 | 0.01 | 0.95 | 17.32 | 0.00 | 0.08 | 0.79 |
| Grooming others | **1.2** ± 1.1 | **0.1** ± 0.1 | 0.0 ± 0.0 | 0.0 ± 0.0 | 2.6 ± 2.2 | 0.2 ± 0.2 | 1.76 | 0.20 | 2.44 | 0.13 | 1.76 | 0.20 |
| Hiding alone | **43.4** ± 15.2 | **57.4** ± 24.6 | 50.1 ± 13.0 | 92.8 ± 39.7 | 35.4 ± 29.3 | 11.8 ± 4.5 | 0.09 | 0.77 | 2.17 | 0.15 | 1.05 | 0.32 |
| Hiding social | **45.5** ± 11.3 | **30.8** ± 8.3 | 68.7 ± 14.9 | 40.5 ± 13.2 | 17.6 ± 3.8 | 18.4 ± 5.7 | 1.16 | 0.29 | 8.25 | 0.01 | 1.29 | 0.27 |
| Any other behavior | **14.0** ± 5.8 | **6.5** ± 2.3 | 0.4 ± 0.4 | 0.2 ± 0.2 | 30.3 ± 8.1 | 14.6 ± 3.3 | 4.31 | 0.05 (0.11) | 33.47 | 0.00 | 4.03 | 0.06 |
| Non-social exploration | **216.6** ± 23.6 | **200.5** ± 12.9 | 254.2 ± 19.2 | 212.2 ± 11.3 | 171.5 ± 37.7 | 185.4 ± 24.5 | 0.33 | 0.57 | 5.03 | 0.04 | 1.31 | 0.26 |
| RestingPassive/ immobile alone | **77.5** ± 7.4 | **99.8** ± 18.0 | 72.8 ± 6.4 | 122.1 ± 27.3 | 83.2 ± 13.9 | 71.2 ± 16.0 | 0.65 | 0.43 | 0.77 | 0.39 | 1.77 | 0.20 |
| RestingPassive/immobile social | **4.6** ± 1.6 | **14.5** ± 4.2 | 4.4 ± 1.7 | 19.3 ± 6.9 | 4.9 ± 2.9 | 8.4 ± 2.5 | 2.81 | 0.11 | 0.91 | 0.35 | 1.11 | 0.30 |
| Rearing supported | **160.3** ± 14.0 | **152.7** ± 13.3 | 196.3 ± 7.2 | 162.6 ± 20.2 | 117.0 ± 13.8 | 139.8 ± 14.6 | 0.09 | 0.77 | 7.72 | 0.01 | 2.37 | 0.14 |
| Rearing unsupported | **28.4** ± 7.2 | **19.9** ± 5.1 | 37.1 ± 10.9 | 19.4 ± 5.1 | 17.8 ± 6.5 | 20.5 ± 9.6 | 0.70 | 0.41 | 1.05 | 0.32 | 1.29 | 0.27 |
| Self grooming | **258.2** ± 59.1 | **316.8** ± 40.2 | 137.3 ± 21.8 | 241.9 ± 54.5 | 403.2 ± 92.3 | 413.1 ± 34.2 | 0.92 | 0.35 | 13.41 | 0.00 | 0.63 | 0.44 |
| Sniffing anogenitally | **246.7** ± 57.9 | **219.6** ± 22.7 | 140.9 ± 14.9 | 174.2 ± 12.2 | 373.6 ± 100.1 | 277.9 ± 39.8 | 0.41 | 0.53 | 12.04 | 0.00 | 1.77 | 0.20 |
| Sniffing body | **104.1** ± 11.2 | **94.8** ± 11.6 | 78.7 ± 11.1 | 77.2 ± 9.7 | 134.7 ± 9.6 | 117.3 ± 20.4 | 0.38 | 0.54 | 9.83 | 0.01 | 0.27 | 0.61 |
| Sniffing nose-to-nose | **303.8** ± 20.6 | **365.8** ± 36.5 | 327.3 ± 24.5 | 411.6 ± 51.5 | 275.7 ± 30.0 | 307.0 ± 41.0 | 1.45 | 0.24 | 2.65 | 0.12 | 0.30 | 0.59 |
| Stand-off/Nose-off | **89.7** ± 31.8 | **61.4** ± 23.9 | 108.2 ± 56.1 | 40.6 ± 12.0 | 67.5 ± 13.4 | 88.1 ± 50.7 | 0.32 | 0.58 | 0.01 | 0.93 | 1.13 | 0.30 |
| Fighting/wrestling | **8.4** ± 2.9 | **4.9** ± 1.4 | 7.7 ± 3.8 | 4.6 ± 1.5 | 9.3 ± 4.4 | 5.3 ± 2.5 | 1.26 | 0.27 | 0.14 | 0.72 | 0.02 | 0.89 |
| Ratio Act/Passve | **27.7** ± 2.4 | **58.9** ± 27.8 | 30.9 ± 2.5 | 35.6 ± 16.1 | 23.9 ± 3.7 | 88.9 ± 58.3 | 0.96 | 0.34 | 0.42 | 0.52 | 0.72 | 0.41 |
| Ratio Social activity | **0.24** ± 0.0 | **0.24** ± 0.0 | 0.21 ± 0.0 | 0.22 ± 0.0 | 0.28 ± 0.0 | 0.26 ± 0.0 | 0.03 | 0.86 | 6.75 | 0.02 | 0.35 | 0.56 |
| Sniffing anogenitally ratio | **0.34** ± 0.0 | **0.33** ± 0.0 | 0.26 ± 0.0 | 0.28 ± 0.0 | 0.44 ± 0.1 | 0.40 ± 0.0 | 0.07 | 0.79 | 15.46 | 0.00 | 0.57 | 0.46 |
| Sniffing body ratio | **0.16** ± 0.0 | **0.14** ± 0.0 | 0.14 ± 0.0 | 0.12 ± 0.0 | 0.18 ± 0.0 | 0.17 ± 0.0 | 1.11 | 0.30 | 5.62 | 0.03 | 0.06 | 0.82 |
| Sniffing nose ratio | **0.50** ± 0.0 | **0.53** ± 0.0 | 0.60 ± 0.0 | 0.60 ± 0.0 | 0.38 ± 0.1 | 0.44 ± 0.0 | 0.51 | 0.48 | 21.12 | 0.00 | 0.31 | 0.58 |
| % unsupported rearing | **12.94** ± 2.7 | **9.19** ± 2.0 | 14.34 ± 4.1 | 8.76 ± 2.2 | 11.25 ± 3.2 | 9.75 ± 3.6 | 1.01 | 0.33 | 0.09 | 0.77 | 0.33 | 0.57 |

Supplementary Table 2

| | | | DEV | CTR | Females DEV | Females CTR | Males DEV | Males CTR | Effect of treatment F | Effect of treatment p* | Effect of sex F | Effect of sex p | Interaction effect F | Interaction effect p |
|---|---|---|---|---|---|---|---|---|---|---|---|---|---|---|
| | Last and first initiated behavior duration | Last 10 avg duration | **2.8** ± 0.2 | **3.1** ± 0.3 | 3.0 ± 0.3 | 3.5 ± 0.5 | 2.5 ± 0.2 | 2.7 ± 0.4 | 0.56 | 0.46 | 2.22 | 0.15 | 0.17 | 0.69 |
| | | First 10 avg duration | **2.9** ± 0.2 | **3.0** ± 0.2 | 2.8 ± 0.3 | 2.8 ± 0.2 | 3.1 ± 0.1 | 3.2 ± 0.3 | 0.03 | 0.88 | 1.30 | 0.27 | 0.05 | 0.82 |
| | Initiator vs responder | Continuation behaviors | **18.8** ± 3.5 | **43.0** ± 19.5 | 12.1 ± 3.7 | 20.7 ± 6.6 | 26.7 ± 3.9 | 71.6 ± 41.4 | 1.20 | 0.28 | 1.81 | 0.19 | 0.56 | 0.46 |
| | | Initiator | **681.4** ± 59.0 | **650.3** ± 47.9 | 614.0 ± 23.4 | 647.0 ± 45.7 | 762.2 ± 116.8 | 654.6 ± 92.3 | 0.22 | 0.65 | 0.95 | 0.34 | 0.77 | 0.39 |
| | | Responder | **53.9** ± 8.9 | **53.2** ± 13.0 | 36.6 ± 8.0 | 40.5 ± 8.3 | 74.5 ± 11.5 | 69.7 ± 26.5 | 0.00 | 0.98 | 3.65 | 0.07 | 0.06 | 0.81 |
| | | Ratio initiator (initiated beh/all beh) | **0.9** ± 0.0 | **0.9** ± 0.0 | 0.9 ± 0.0 | 0.9 ± 0.0 | 0.9 ± 0.0 | 0.8 ± 0.1 | 0.35 | 0.56 | 2.99 | 0.10 | 0.10 | 0.75 |
| Duration | Initiator | Initiated social investigation | **620.6** ± 63.3 | **629.9** ± 49.9 | 524.2 ± 29.8 | 621.0 ± 52.8 | 736.2 ± 115.0 | 641.3 ± 91.5 | 0.00 | 0.99 | 2.00 | 0.17 | 1.36 | 0.26 |
| | | Initiated fighting | **4.8** ± 2.2 | **1.8** ± 0.7 | 6.5 ± 3.8 | 2.1 ± 1.0 | 2.8 ± 1.4 | 1.5 ± 0.9 | 1.80 | 0.19 | 1.00 | 0.33 | 0.50 | 0.49 |
| | | Initiated grooming others | **0.1** ± 0.1 | **0.1** ± 0.1 | 0.0 ± 0.0 | 0.0 ± 0.0 | 0.1 ± 0.1 | 0.2 ± 0.2 | 0.09 | 0.76 | 1.99 | 0.17 | 0.09 | 0.76 |
| | | Initiated Sniffing anogenitally | **238.5** ± 55.1 | **208.9** ± 22.1 | 138.9 ± 13.8 | 171.9 ± 11.7 | 358.0 ± 96.0 | 256.4 ± 41.8 | 0.52 | 0.48 | 10.24 | 0.00 | 2.01 | 0.17 |
| | | Initiated Sniffing body | **101.1** ± 11.2 | **88.4** ± 11.2 | 75.9 ± 10.3 | 72.9 ± 10.3 | 131.2 ± 10.9 | 108.4 ± 19.6 | 0.73 | 0.40 | 8.94 | 0.01 | 0.42 | 0.52 |
| | | Initiated Sniffing nose | **281.0** ± 20.7 | **332.6** ± 34.9 | 309.4 ± 22.7 | 376.2 ± 47.9 | 247.0 ± 30.0 | 276.5 ± 42.1 | 1.10 | 0.31 | 3.10 | 0.09 | 0.17 | 0.69 |
| | | Init Stand-off | **55.9** ± 27.3 | **18.5** ± 6.1 | 83.3 ± 46.9 | 23.9 ± 10.1 | 23.0 ± 5.9 | 11.6 ± 4.0 | 2.08 | 0.16 | 2.20 | 0.15 | 0.96 | 0.34 |
| | | Initiated behaviors | **681.4** ± 59.0 | **650.3** ± 47.9 | 614.0 ± 23.4 | 647.0 ± 45.7 | 762.2 ± 116.8 | 654.6 ± 92.3 | 0.22 | 0.65 | 0.95 | 0.34 | 0.77 | 0.39 |
| | Responder | Responded social invest | **23.0** ± 4.8 | **25.9** ± 4.0 | 19.1 ± 5.0 | 25.6 ± 5.5 | 27.7 ± 8.2 | 26.3 ± 5.8 | 0.14 | 0.71 | 0.47 | 0.50 | 0.35 | 0.56 |
| | | Resp Fighting | **3.6** ± 1.7 | **2.3** ± 0.8 | 1.2 ± 0.8 | 2.3 ± 1.2 | 6.5 ± 3.2 | 2.4 ± 1.1 | 0.71 | 0.41 | 2.41 | 0.14 | 2.13 | 0.16 |
| | | Resp Sniffing anog | **0.1** ± 0.1 | **0.5** ± 0.3 | 0.0 ± 0.0 | 0.3 ± 0.2 | 0.2 ± 0.2 | 0.7 ± 0.6 | 1.03 | 0.32 | 0.46 | 0.50 | 0.03 | 0.87 |
| | | Resp Sniffing body | **2.2** ± 0.9 | **1.8** ± 0.7 | 2.3 ± 1.3 | 0.4 ± 0.4 | 2.0 ± 1.1 | 3.7 ± 1.2 | 0.01 | 0.92 | 2.10 | 0.16 | 2.86 | 0.11 |
| | | Resp Sniffing nose | **20.7** ± 4.4 | **23.6** ± 3.7 | 16.8 ± 4.2 | 24.9 ± 5.3 | 25.4 ± 7.6 | 21.9 ± 5.0 | 0.14 | 0.72 | 0.21 | 0.65 | 0.89 | 0.36 |
| | | Resp Stand-off | **27.3** ± 7.1 | **25.0** ± 11.2 | 16.3 ± 7.4 | 12.6 ± 3.8 | 40.4 ± 10.2 | 40.9 ± 23.8 | 0.01 | 0.92 | 3.05 | 0.09 | 0.02 | 0.89 |
| | | Behaviors responded | **53.9** ± 8.9 | **53.2** ± 13.0 | 36.6 ± 8.0 | 40.5 ± 8.3 | 74.5 ± 11.5 | 69.7 ± 26.5 | 0.00 | 0.98 | 3.65 | 0.07 | 0.06 | 0.81 |

Supplementary Table 2

| | | | DEV | CTR | Females DEV | Females CTR | Males DEV | Males CTR | Effect of treatment F | Effect of treatment p* | Effect of sex F | Effect of sex p | Interaction effect F | Interaction effect p |
|---|---|---|---|---|---|---|---|---|---|---|---|---|---|---|
| **Duration** | **Tactile contact** | Total time in contact | **872.7** ± 64.6 | **887.3** ± 47.4 | 821.9 ± 60.9 | 880.0 ± 55.7 | 933.7 ± 116.2 | 896.8 ± 81.2 | 0.02 | 0.90 | 0.59 | 0.45 | 0.32 | 0.58 |
| | | Average time in contact | **2.9** ± 0.1 | **3.2** ± 0.1 | 3.1 ± 0.2 | 3.3 ± 0.1 | 2.8 ± 0.2 | 3.1 ± 0.2 | 1.38 | 0.25 | 1.48 | 0.24 | 0.01 | 0.92 |
| | | Average length of first 10 behaviors with tactile contact | **2.8** ± 0.2 | **3.3** ± 0.2 | 2.8 ± 0.4 | 3.0 ± 0.3 | 2.7 ± 0.3 | 3.6 ± 0.3 | 2.30 | 0.14 | 0.46 | 0.51 | 0.78 | 0.39 |
| | | Average length of last 10 behaviors with tactile contact | **2.9** ± 0.2 | **3.4** ± 0.3 | 2.8 ± 0.3 | 3.9 ± 0.4 | 3.0 ± 0.3 | 2.9 ± 0.4 | 1.54 | 0.23 | 0.66 | 0.43 | 1.96 | 0.18 |
| | | Not in contact(OA&OA_near_walls) | **1294.0** ± 126.5 | **1170.7** ± 105.3 | 1071.7 ± 89.3 | 929.6 ± 118.0 | 1560.9 ± 199.8 | 1480.6 ± 102.9 | 0.63 | 0.44 | 13.69 | 0.00 | 0.05 | 0.83 |
| | | Beh-wo-contact(OA&OA_near_walls) | **163.9** ± 22.3 | **140.0** ± 16.2 | 136.8 ± 23.6 | 106.4 ± 18.6 | 196.4 ± 35.0 | 183.3 ± 18.0 | 0.74 | 0.40 | 7.32 | 0.01 | 0.12 | 0.74 |
| | | Time in OA without social interaction | **1457.9** ± 140.3 | **1310.7** ± 117.8 | 1208.4 ± 98.1 | 1035.9 ± 134.5 | 1757.3 ± 220.4 | 1663.9 ± 104.7 | 0.74 | 0.40 | 14.40 | 0.00 | 0.07 | 0.80 |
| **Duration** | **Conflict towards CTR/DEV** | Fighting/boxing/wrestling /kicking towards DEV | **3.3** ± 1.8 | **4.2** ± 1.3 | 3.4 ± 2.4 | 4.0 ± 1.4 | 3.2 ± 2.9 | 4.4 ± 2.5 | 0.13 | 0.72 | 0.00 | 0.97 | 0.02 | 0.90 |
| | | Stand-off/Nose-off towards DEV | **25.0** ± 7.5 | **38.0** ± 15.7 | 21.1 ± 11.5 | 21.0 ± 6.5 | 29.6 ± 8.4 | 59.9 ± 33.1 | 0.53 | 0.48 | 1.31 | 0.26 | 0.54 | 0.47 |
| | | Conflict towards DEV | **28.3** ± 8.9 | **42.2** ± 16.4 | 24.5 ± 13.8 | 25.0 ± 5.9 | 32.8 ± 10.0 | 64.3 ± 35.0 | 0.52 | 0.48 | 1.17 | 0.29 | 0.49 | 0.49 |
| | | Adjusted Conflict towards DEV | **11.2** ± 2.8 | **14.4** ± 5.4 | 9.4 ± 4.4 | 8.3 ± 1.7 | 13.3 ± 2.8 | 22.3 ± 11.6 | 0.31 | 0.59 | 1.55 | 0.23 | 0.50 | 0.49 |
| | | Fighting/boxing/wrestling /kicking towards CTR | **5.1** ± 2.1 | **0.7** ± 0.4 | 4.3 ± 3.1 | 0.6 ± 0.4 | 6.1 ± 2.8 | 0.9 ± 0.7 | 4.99 | 0.04 (0.1) | 0.30 | 0.59 | 0.14 | 0.71 |
| | | Stand-off/Nose-off towards CTR | **64.7** ± 31.6 | **17.7** ± 4.8 | 87.1 ± 56.0 | 19.6 ± 7.1 | 37.9 ± 7.2 | 15.3 ± 5.9 | 2.50 | 0.13 | 0.88 | 0.36 | 0.62 | 0.44 |
| | | Conflic towards CTR | **69.8** ± 31.3 | **18.5** ± 4.9 | 91.3 ± 55.3 | 20.2 ± 7.3 | 44.0 ± 10.0 | 16.2 ± 6.1 | 3.06 | 0.09 | 0.82 | 0.37 | 0.59 | 0.45 |
| | | Adjusted Conflic towards CTR | **18.2** ± 7.8 | **6.3** ± 1.6 | 23.1 ± 13.8 | 6.8 ± 2.4 | 12.3 ± 2.8 | 5.6 ± 2.1 | 2.56 | 0.12 | 0.70 | 0.41 | 0.45 | 0.51 |

Supplementary Table 2

| | | | | Females | | Males | | Effect of treatment | | Effect of sex | | Interaction effect | |
|---|---|---|---|---|---|---|---|---|---|---|---|---|---|
| | | **DEV** | **CTR** | DEV | CTR | DEV | CTR | F | p* | F | p | F | p |
| Duration – Following | DEV | **19.1** ± 5.9 | **25.3** ± 7.7 | 5.2 ± 2.4 | 6.9 ± 1.9 | 35.6 ± 7.8 | 46.4 ± 12.8 | 0.53 | 0.47 | 16.96 | 0.00 | 0.28 | 0.60 |
| | Adjusted following DEV | **9.7** ± 3.0 | **9.8** ± 3.7 | 2.9 ± 1.3 | 2.4 ± 0.7 | 17.7 ± 4.1 | 19.2 ± 7.0 | 0.01 | 0.91 | 12.78 | 0.00 | 0.05 | 0.82 |
| | CTR | **36.8** ± 18.9 | **33.3** ± 11.9 | 12.1 ± 3.9 | 7.5 ± 2.5 | 66.5 ± 37.1 | 62.6 ± 21.4 | 0.04 | 0.84 | 7.17 | 0.01 | 0.00 | 0.99 |
| | Adjusted following CTR | **9.3** ± 4.7 | **9.7** ± 3.8 | 3.0 ± 1.0 | 2.3 ± 0.8 | 17.0 ± 9.2 | 19.2 ± 7.1 | 0.02 | 0.89 | 7.57 | 0.01 | 0.07 | 0.79 |
| Duration – Interaction bout length | Interaction bout length w DEV | **3.0** ± 0.1 | **3.0** ± 0.1 | 2.8 ± 0.2 | 2.9 ± 0.1 | 3.1 ± 0.2 | 3.1 ± 0.2 | 0.00 | 0.98 | 1.90 | 0.18 | 0.02 | 0.89 |
| | Interaction bout length w CTR | **3.1** ± 0.3 | **3.3** ± 0.1 | 3.4 ± 0.4 | 3.4 ± 0.2 | 2.7 ± 0.2 | 3.1 ± 0.1 | 0.81 | 0.40 | 2.78 | 0.11 | 0.49 | 0.49 |
| | Interaction bout length DD & CC | **3.0** ± 0.1 | **3.3** ± 0.1 | 2.8 ± 0.2 | 3.4 ± 0.2 | 3.1 ± 0.2 | 3.1 ± 0.1 | 1.92 | 0.18 | 0.01 | 0.93 | 1.83 | 0.19 |
| Duration – Receiving behaviors | Receiveing all behaviors | **804.2** ± 70.1 | **787.7** ± 37.9 | 972.8 ± 71.4 | 855.1 ± 41.5 | 601.9 ± 37.8 | 700.9 ± 52.6 | 0.03 | 0.87 | 20.68 | 0.00 | 3.52 | 0.07 |
| | Receiveing Active social | **652.2** ± 60.9 | **655.8** ± 47.6 | 768.4 ± 80.4 | 721.7 ± 65.2 | 512.8 ± 38.9 | 571.1 ± 54.6 | 0.01 | 0.94 | 8.04 | 0.01 | 0.54 | 0.47 |
| | Receiveing anogenital sniffing | **250.3** ± 38.8 | **200.1** ± 22.0 | 337.7 ± 45.2 | 238.8 ± 31.8 | 145.5 ± 17.5 | 150.4 ± 14.8 | 1.88 | 0.18 | 16.78 | 0.00 | 2.29 | 0.14 |
| | Receiveing Sniffing body | **93.7** ± 10.4 | **95.8** ± 9.8 | 97.5 ± 16.5 | 95.3 ± 16.5 | 89.1 ± 11.1 | 96.5 ± 7.5 | 0.03 | 0.87 | 0.05 | 0.82 | 0.09 | 0.77 |
| | Receiveing Sniffing Nose | **308.3** ± 25.8 | **359.8** ± 36.9 | 333.2 ± 38.6 | 387.5 ± 52.7 | 278.3 ± 27.4 | 324.3 ± 46.9 | 0.92 | 0.35 | 1.28 | 0.27 | 0.01 | 0.94 |
| | Being Followed | **52.9** ± 16.1 | **56.5** ± 17.4 | 82.9 ± 22.8 | 75.5 ± 28.2 | 17.0 ± 6.3 | 32.0 ± 10.4 | 0.02 | 0.88 | 4.96 | 0.04 | 0.21 | 0.65 |
| | Receiveing Conflict | **89.6** ± 25.3 | **66.5** ± 23.1 | 104.9 ± 43.3 | 45.9 ± 10.6 | 71.3 ± 16.9 | 92.8 ± 49.3 | 0.26 | 0.62 | 0.03 | 0.86 | 1.18 | 0.29 |
| Duration – Burrow (Tunnels + nestboxes) | Social investigation | **350.41** ± 45.54 | **406.94** ± 44.51 | 372.6 ± 23.0 | 490.2 ± 55.4 | 323.8 ± 94.95 | 299.9 ± 48.69 | 0.53 | 0.47 | 3.48 | 0.08 | 1.22 | 0.28 |
| | Non-social invest | **912.37** ± 119.64 | **960.97** ± 88.47 | 1179.5 ± 78.1 | 1158.9 ± 91.6 | 591.8 ± 151.11 | 706.5 ± 102.80 | 0.17 | 0.69 | 20.16 | 0.00 | 0.34 | 0.57 |
| | Interacting with the environment | **20.2** ± 7.7 | **30.1** ± 10.5 | 26.4 ± 10.8 | 41.2 ± 17.0 | 12.8 ± 9.8 | 15.8 ± 6.8 | 0.37 | 0.55 | 1.75 | 0.20 | 0.16 | 0.69 |
| | Fighting/boxing/wrestling /kicking | **5.0** ± 2.4 | **1.8** ± 0.8 | 5.5 ± 3.3 | 2.6 ± 1.2 | 4.4 ± 3.5 | 0.8 ± 0.6 | 1.85 | 0.19 | 0.35 | 0.56 | 0.02 | 0.88 |
| | Following | **14.6** ± 4.1 | **12.8** ± 4.3 | 9.3 ± 2.8 | 7.8 ± 3.0 | 21.0 ± 7.4 | 19.2 ± 8.3 | 0.07 | 0.79 | 3.47 | 0.08 | 0.00 | 0.98 |
| | Any other behavior | **1.3** ± 0.9 | **0.4** ± 0.3 | 0.0 ± 0.0 | 0.0 ± 0.0 | 2.8 ± 1.7 | 0.9 ± 0.6 | 1.59 | 0.22 | 5.84 | 0.02 | 1.59 | 0.22 |
| | Non-social exploration | **99.5** ± 21.8 | **87.7** ± 12.7 | 155.3 ± 19.2 | 114.9 ± 12.2 | 32.5 ± 10.7 | 52.7 ± 17.0 | 0.36 | 0.56 | 29.98 | 0.00 | 3.22 | 0.09 |
| | Passive/ immobile alone | **48.7** ± 9.4 | **87.6** ± 18.8 | 68.7 ± 6.7 | 119.6 ± 28.0 | 24.7 ± 12.3 | 46.5 ± 11.1 | 2.62 | 0.12 | 6.79 | 0.02 | 0.42 | 0.53 |
| | Passive/immobile social | **2.4** ± 1.1 | **12.6** ± 4.1 | 2.9 ± 1.6 | 18.0 ± 6.3 | 1.7 ± 1.5 | 5.7 ± 2.9 | 3.56 | 0.07 | 1.81 | 0.19 | 1.19 | 0.29 |

Supplementary Table 2

| | | | | Females | | Males | | Effect of treatment | | Effect of sex | | Interaction effect | |
|---|---|---|---|---|---|---|---|---|---|---|---|---|---|
| | | **DEV** | **CTR** | DEV | CTR | DEV | CTR | F | p* | F | p | F | p |
| Duration – Burrow (Tunnels + nestboxes) | Rearing supported | **20.0** ± 6.5 | **13.1** ± 5.0 | 33.5 ± 8.5 | 23.4 ± 7.2 | 3.8 ± 2.1 | 0.0 ± 0.0 | 1.05 | 0.32 | 15.06 | 0.00 | 0.21 | 0.65 |
| | Rearing unsupported | **1.0** ± 0.9 | **0.9** ± 0.5 | 1.8 ± 1.6 | 1.6 ± 0.8 | 0.0 ± 0.0 | 0.0 ± 0.0 | 0.01 | 0.92 | 2.84 | 0.11 | 0.01 | 0.92 |
| | Self grooming | **61.4** ± 15.2 | **143.9** ± 38.5 | 95.2 ± 17.8 | 197.5 ± 56.2 | 20.9 ± 7.5 | 74.9 ± 36.1 | 2.73 | 0.11 | 4.33 | 0.05 | 0.26 | 0.62 |
| | Sniffing anogenitally | **124.0** ± 33.1 | **116.1** ± 11.3 | 87.9 ± 12.1 | 124.8 ± 14.1 | 167.4 ± 66.3 | 105.0 ± 17.6 | 0.17 | 0.69 | 0.90 | 0.35 | 2.49 | 0.13 |
| | Sniffing body | **33.2** ± 6.2 | **30.7** ± 5.0 | 41.5 ± 8.7 | 41.2 ± 6.7 | 23.3 ± 6.2 | 17.1 ± 3.4 | 0.19 | 0.67 | 8.39 | 0.01 | 0.16 | 0.69 |
| | Sniffing nose-to-nose | **193.1** ± 23.3 | **260.1** ± 34.4 | 243.2 ± 17.8 | 324.2 ± 45.6 | 133.1 ± 29.0 | 177.7 ± 32.3 | 2.33 | 0.14 | 9.69 | 0.01 | 0.20 | 0.66 |
| | All sniffing | ± 45.5 | ± 44.5 | ± 23.0 | ± 55.4 | ± 95.0 | ± 48.7 | 0.53 | 0.47 | 3.48 | 0.08 | 1.22 | 0.28 |
| | Stand-off/Nose-off | **73.0** ± 32.5 | **50.0** ± 23.1 | 98.0 ± 56.7 | 32.9 ± 12.4 | 43.0 ± 12.4 | 72.0 ± 49.1 | 0.19 | 0.66 | 0.04 | 0.85 | 1.31 | 0.26 |
| | Walking/running | **812.9** ± 104.7 | **873.3** ± 78.8 | 1024.2 ± 78.3 | 1044.0 ± 86.1 | 559.4 ± 143.8 | 653.8 ± 89.4 | 0.28 | 0.60 | 15.67 | 0.00 | 0.12 | 0.73 |
| | Burrow Total | **1510.4** ± 190.4 | **1721.1** ± 163.9 | 1893.4 ± 131.1 | 2093.6 ± 192.9 | 1050.7 ± 270.7 | 1242.2 ± 143.5 | 0.88 | 0.36 | 16.43 | 0.00 | 0.00 | 0.98 |
| Duration – Full OA (OA+OA near walls) | Social investigation | **289.8** ± 47.3 | **261.4** ± 40.3 | 157.7 ± 16.1 | 161.4 ± 26.9 | 448.2 ± 35.2 | 390.0 ± 55.8 | 0.44 | 0.51 | 39.77 | 0.00 | 0.57 | 0.46 |
| | Non-social invest | **1186.1** ± 81.7 | **1070.7** ± 83.4 | 1104.7 ± 98.5 | 935.9 ± 118.1 | 1283.7 ± 122.0 | 1244.0 ± 75.3 | 0.76 | 0.39 | 4.13 | 0.05 | 0.29 | 0.60 |
| | Interacting with the environment | **11.7** ± 4.9 | **12.7** ± 3.0 | 16.6 ± 8.3 | 14.8 ± 4.8 | 5.8 ± 2.2 | 9.9 ± 2.5 | 0.04 | 0.84 | 1.91 | 0.18 | 0.27 | 0.61 |
| | Fighting/boxing/wrestling /kicking | **3.4** ± 1.5 | **3.1** ± 1.2 | 2.2 ± 1.4 | 2.0 ± 1.0 | 4.9 ± 2.8 | 4.5 ± 2.3 | 0.02 | 0.88 | 1.61 | 0.22 | 0.00 | 0.97 |
| | Following | **41.3** ± 17.5 | **42.4** ± 14.3 | 8.1 ± 4.1 | 5.1 ± 1.7 | 81.1 ± 29.7 | 90.3 ± 21.9 | 0.03 | 0.86 | 19.92 | 0.00 | 0.12 | 0.73 |
| | Grooming others | **1.2** ± 1.1 | **0.1** ± 0.1 | 0.0 ± 0.0 | 0.0 ± 0.0 | 2.6 ± 2.2 | 0.2 ± 0.2 | 1.76 | 0.20 | 2.43 | 0.13 | 1.76 | 0.20 |
| | Hiding alone | **43.4** ± 15.2 | **57.4** ± 24.6 | 50.1 ± 13.0 | 92.8 ± 39.7 | 35.4 ± 29.3 | 11.8 ± 4.5 | 0.09 | 0.77 | 2.17 | 0.15 | 1.05 | 0.32 |
| | Hiding social | **45.5** ± 11.3 | **30.8** ± 8.3 | 68.7 ± 14.9 | 40.5 ± 13.2 | 17.6 ± 3.8 | 18.4 ± 5.7 | 1.16 | 0.29 | 8.25 | 0.01 | 1.29 | 0.27 |
| | Any other behavior | **12.7** ± 5.7 | **6.1** ± 2.1 | 0.4 ± 0.4 | 0.2 ± 0.2 | 27.5 ± 8.7 | 13.7 ± 3.0 | 3.07 | 0.09 | 25.92 | 0.00 | 2.85 | 0.11 |
| | Non-social exploration | **96.3** ± 13.8 | **96.4** ± 13.3 | 81.4 ± 12.2 | 71.6 ± 11.6 | 114.2 ± 24.4 | 128.2 ± 21.0 | 0.01 | 0.91 | 5.62 | 0.03 | 0.40 | 0.53 |
| | Passive/ immobile alone | **28.4** ± 12.1 | **12.2** ± 4.2 | 3.3 ± 1.4 | 2.5 ± 1.6 | 58.5 ± 19.2 | 24.6 ± 6.8 | 3.78 | 0.06 | 18.81 | 0.00 | 3.46 | 0.08 |
| | Passive/immobile social | **2.3** ± 1.4 | **1.9** ± 0.7 | 1.4 ± 0.9 | 1.3 ± 0.9 | 3.3 ± 2.9 | 2.7 ± 1.2 | 0.06 | 0.82 | 0.99 | 0.33 | 0.02 | 0.88 |
| | Rearing supported | **140.2** ± 11.0 | **139.5** ± 11.8 | 162.8 ± 10.3 | 139.3 ± 17.6 | 113.1 ± 13.0 | 139.8 ± 14.6 | 0.01 | 0.93 | 2.05 | 0.17 | 2.14 | 0.16 |
| | Rearing unsupported | **27.4** ± 6.9 | **18.7** ± 5.0 | 35.4 ± 10.3 | 17.4 ± 4.8 | 17.8 ± 6.5 | 20.5 ± 9.6 | 0.77 | 0.39 | 0.69 | 0.42 | 1.41 | 0.25 |
| | Self grooming | **196.8** ± 68.4 | **172.9** ± 40.3 | 42.1 ± 8.2 | 44.4 ± 11.8 | 382.4 ± 99.6 | 338.2 ± 36.3 | 0.20 | 0.66 | 45.64 | 0.00 | 0.25 | 0.63 |
| | Sniffing anogenitally | **121.2** ± 29.1 | **102.3** ± 19.7 | 51.8 ± 7.3 | 48.8 ± 8.1 | 204.4 ± 38.4 | 171.1 ± 26.8 | 0.62 | 0.44 | 35.34 | 0.00 | 0.43 | 0.52 |
| | Sniffing body | **67.9** ± 11.8 | **61.4** ± 12.0 | 34.2 ± 7.0 | 33.7 ± 5.9 | 108.5 ± 2.5 | 97.1 ± 19.2 | 0.23 | 0.64 | 30.81 | 0.00 | 0.20 | 0.66 |
| | Sniffing nose-to-nose | **99.5** ± 17.6 | **97.6** ± 12.3 | 71.7 ± 6.9 | 78.9 ± 16.9 | 132.7 ± 32.1 | 121.7 ± 13.0 | 0.01 | 0.92 | 6.78 | 0.02 | 0.21 | 0.65 |
| | All sniffing | **288.6** ± 46.8 | **261.3** ± 40.3 | 157.7 ± 16.1 | 161.4 ± 26.9 | 445.6 ± 34.1 | 389.8 ± 55.8 | 0.40 | 0.53 | 39.63 | 0.00 | 0.53 | 0.48 |
| | Stand-off/Nose-off | **16.2** ± 3.9 | **11.2** ± 2.5 | 9.2 ± 2.5 | 7.6 ± 2.4 | 24.6 ± 6.2 | 15.8 ± 4.1 | 1.63 | 0.22 | 8.32 | 0.01 | 0.78 | 0.39 |
| | Walking/running | **1089.7** ± 74.9 | **974.3** ± 75.5 | 1023.2 ± 89.2 | 864.2 ± 108.6 | 1169.5 ± 115.7 | 1115.8 ± 72.1 | 0.92 | 0.35 | 3.21 | 0.09 | 0.22 | 0.64 |
| | Full OA Total | **2045.3** ± 187.1 | **1841.0** ± 164.1 | 1663.1 ± 129.8 | 1465.2 ± 192.3 | 2504.0 ± 260.9 | 2324.3 ± 142.4 | 0.84 | 0.37 | 17.01 | 0.00 | 0.00 | 0.97 |

*in parenthesis, p corrected with B-H procedure where applicable

CTR - Control, DEV - Devocalized

Supplementary Table 3

Nr of instances

Behavior in the whole environment

| | | | Females | | Males | | Effect of treatment | | Effect of sex | | Interaction effect | |
|---|---|---|---|---|---|---|---|---|---|---|---|---|
| | **DEV** | **CTR** | DEV | CTR | DEV | CTR | F | p* | F | p | F | p |
| Social investigation | **237.8** ± 19.0 | **227.8** ± 19.7 | 197.8 ± 17.8 | 212.9 ± 18.7 | 285.8 ± 21.3 | 246.9 ± 36.9 | 0.18 | 0.68 | 4.67 | 0.04 | 0.92 | 0.35 |
| Non-social invest | **520.9** ± 29.7 | **491.1** ± 24.0 | 558.8 ± 46.9 | 508.1 ± 33.7 | 475.4 ± 18.1 | 469.3 ± 32.0 | 0.52 | 0.48 | 2.41 | 0.13 | 0.32 | 0.58 |
| Immobile+hiding | **29.3** ± 4.1 | **26.9** ± 3.9 | 38.2 ± 4.7 | 33.9 ± 5.7 | 18.6 ± 2.5 | 17.9 ± 2.6 | 0.24 | 0.63 | 12.06 | 0.00 | 0.12 | 0.73 |
| All Rearing | **66.2** ± 7.1 | **61.0** ± 6.9 | 80.8 ± 7.3 | 63.9 ± 9.9 | 48.6 ± 7.4 | 57.3 ± 9.0 | 0.17 | 0.69 | 3.68 | 0.07 | 1.60 | 0.22 |
| Conflict | **17.7** ± 3.0 | **13.1** ± 2.5 | 16.2 ± 3.8 | 11.6 ± 2.6 | 19.6 ± 4.7 | 15.1 ± 4.5 | 1.17 | 0.29 | 0.70 | 0.41 | 0.00 | 0.99 |
| All Sniffing | **237.6** ± 19.0 | **227.7** ± 19.7 | 197.8 ± 17.8 | 212.9 ± 18.7 | 285.4 ± 21.2 | 246.7 ± 36.9 | 0.18 | 0.68 | 4.63 | 0.04 | 0.91 | 0.35 |
| All immobile | **11.4** ± 0.7 | **12.5** ± 1.6 | 11.5 ± 0.8 | 13.3 ± 2.1 | 11.2 ± 1.3 | 11.4 ± 2.5 | 0.22 | 0.65 | 0.25 | 0.62 | 0.13 | 0.72 |
| All hiding | **17.9** ± 3.9 | **14.4** ± 3.5 | 26.7 ± 4.7 | 20.6 ± 5.3 | 7.4 ± 1.4 | 6.4 ± 1.7 | 0.57 | 0.46 | 12.67 | 0.00 | 0.30 | 0.59 |
| Non-soc Passive | **18.0** ± 2.2 | **18.4** ± 2.8 | 22.0 ± 2.7 | 23.1 ± 3.9 | 13.2 ± 1.8 | 12.3 ± 2.2 | 0.00 | 0.98 | 7.73 | 0.01 | 0.08 | 0.78 |
| Social Passive (resting+hiding) | **11.3** ± 2.6 | **8.5** ± 1.5 | 16.2 ± 3.7 | 10.8 ± 2.3 | 5.4 ± 1.1 | 5.6 ± 1.1 | 0.98 | 0.33 | 9.15 | 0.01 | 1.11 | 0.30 |
| Walking/running | **471.6** ± 25.7 | **440.4** ± 20.1 | 498.0 ± 40.9 | 452.8 ± 29.1 | 440.0 ± 20.8 | 424.4 ± 25.6 | 0.80 | 0.38 | 1.61 | 0.22 | 0.19 | 0.67 |
| Interacting with the environment | **8.5** ± 2.9 | **7.1** ± 1.1 | 11.7 ± 4.6 | 7.8 ± 1.5 | 4.6 ± 2.2 | 6.3 ± 1.5 | 0.15 | 0.70 | 2.26 | 0.15 | 0.96 | 0.34 |
| Fighting/boxing/wrestling /kicking | **2.3** ± 0.9 | **1.9** ± 0.6 | 2.0 ± 1.3 | 2.1 ± 0.8 | 2.6 ± 1.2 | 1.7 ± 0.9 | 0.12 | 0.73 | 0.01 | 0.93 | 0.20 | 0.66 |
| Following | **15.3** ± 5.5 | **13.9** ± 4.2 | 5.8 ± 2.1 | 3.7 ± 1.0 | 26.6 ± 9.6 | 27.0 ± 6.7 | 0.02 | 0.88 | 15.15 | 0.00 | 0.05 | 0.82 |
| Grooming others | **0.2** ± 0.1 | **0.1** ± 0.1 | 0.0 ± 0.0 | 0.0 ± 0.0 | 0.4 ± 0.2 | 0.1 ± 0.1 | 1.19 | 0.29 | 5.31 | 0.03 | 1.19 | 0.29 |
| Hiding alone | **7.5** ± 1.9 | **8.4** ± 2.7 | 11.7 ± 2.2 | 12.8 ± 4.1 | 2.6 ± 1.2 | 2.7 ± 0.9 | 0.03 | 0.86 | 8.37 | 0.01 | 0.02 | 0.88 |
| Hiding social | **10.4** ± 2.6 | **6.0** ± 1.2 | 15.0 ± 3.7 | 7.8 ± 1.8 | 4.8 ± 1.2 | 3.7 ± 1.1 | 3.05 | 0.09 | 8.98 | 0.01 | 1.66 | 0.21 |
| Any other behavior | **6.6** ± 3.2 | **3.3** ± 1.1 | 0.2 ± 0.2 | 0.1 ± 0.1 | 14.4 ± 5.3 | 7.3 ± 1.6 | 2.33 | 0.14 | 20.81 | 0.00 | 2.26 | 0.15 |
| Non-social exploration | **49.3** ± 5.5 | **50.8** ± 4.8 | 60.8 ± 6.5 | 55.3 ± 5.7 | 35.4 ± 3.6 | 44.9 ± 7.7 | 0.08 | 0.78 | 6.44 | 0.02 | 1.12 | 0.30 |
| Passive/ immobile alone | **10.5** ± 0.8 | **10.0** ± 1.4 | 10.3 ± 0.8 | 10.3 ± 1.9 | 10.6 ± 1.5 | 9.6 ± 2.1 | 0.07 | 0.80 | 0.02 | 0.90 | 0.07 | 0.80 |
| Passive/immobile social | **0.9** ± 0.2 | **2.5** ± 0.5 | 1.2 ± 0.3 | 3.0 ± 0.7 | 0.6 ± 0.4 | 1.9 ± 0.4 | 6.22 | .020 (.1) | 1.90 | 0.18 | 0.22 | 0.65 |
| Rearing supported | **55.2** ± 5.1 | **52.4** ± 5.0 | 67.3 ± 4.1 | 55.8 ± 8.0 | 40.6 ± 4.7 | 48.0 ± 4.3 | 0.09 | 0.77 | 6.04 | 0.02 | 1.82 | 0.19 |
| Rearing unsupported | **11.0** ± 2.6 | **8.6** ± 2.5 | 13.5 ± 3.9 | 8.1 ± 2.1 | 8.0 ± 2.8 | 9.3 ± 5.0 | 0.27 | 0.61 | 0.30 | 0.59 | 0.70 | 0.41 |
| Self grooming | **35.7** ± 5.6 | **37.6** ± 3.1 | 26.5 ± 6.2 | 29.9 ± 3.7 | 46.8 ± 7.3 | 47.6 ± 1.5 | 0.16 | 0.69 | 13.68 | 0.00 | 0.07 | 0.80 |
| Sniffing anogenitally | **73.1** ± 12.1 | **66.9** ± 8.1 | 48.0 ± 5.4 | 55.9 ± 3.3 | 103.2 ± 18.3 | 81.1 ± 16.6 | 0.31 | 0.58 | 10.04 | 0.00 | 1.39 | 0.25 |
| Sniffing body | **45.0** ± 5.0 | **41.8** ± 5.5 | 31.8 ± 4.1 | 34.7 ± 4.2 | 60.8 ± 2.3 | 51.0 ± 10.4 | 0.24 | 0.63 | 10.23 | 0.00 | 0.80 | 0.38 |
| Sniffing nose-to-nose | **119.5** ± 8.0 | **118.9** ± 8.9 | 118.0 ± 10.7 | 122.3 ± 12.8 | 121.4 ± 12.1 | 114.6 ± 11.8 | 0.01 | 0.93 | 0.03 | 0.88 | 0.17 | 0.69 |
| Stand-off/Nose-off | **15.5** ± 2.7 | **11.2** ± 2.2 | 14.2 ± 3.8 | 9.4 ± 2.4 | 17.0 ± 3.6 | 13.4 ± 3.7 | 1.29 | 0.27 | 0.87 | 0.36 | 0.03 | 0.88 |
| Ratio Act/Passve | **48.7** ± 4.9 | **77.8** ± 24.2 | 51.1 ± 6.9 | 75.6 ± 30.5 | 45.8 ± 6.5 | 80.6 ± 38.9 | 0.85 | 0.37 | 0.00 | 1.00 | 0.03 | 0.87 |
| Ratio Social activity | **0.3** ± 0.0 | **0.3** ± 0.0 | 0.3 ± 0.0 | 0.3 ± 0.0 | 0.4 ± 0.0 | 0.3 ± 0.0 | 0.34 | 0.57 | 21.92 | 0.00 | 3.35 | 0.08 |
| Sniffing anogenitally ratio | **0.3** ± 0.0 | **0.3** ± 0.0 | 0.2 ± 0.0 | 0.3 ± 0.0 | 0.3 ± 0.0 | 0.3 ± 0.0 | 0.02 | 0.88 | 10.56 | 0.00 | 1.78 | 0.20 |
| Sniffing body ratio | **0.2** ± 0.0 | **0.2** ± 0.0 | 0.2 ± 0.0 | 0.2 ± 0.0 | 0.2 ± 0.0 | 0.2 ± 0.0 | 0.13 | 0.73 | 8.29 | 0.01 | 0.30 | 0.59 |
| Sniffing nose ratio | **0.5** ± 0.0 | **0.5** ± 0.0 | 0.6 ± 0.0 | 0.6 ± 0.0 | 0.4 ± 0.0 | 0.5 ± 0.0 | 0.18 | 0.68 | 20.87 | 0.00 | 2.18 | 0.15 |
| Overall nr of behaviors | **938.1** ± 44.9 | **881.8** ± 41.1 | 936.2 ± 80.1 | 871.8 ± 52.3 | 940.4 ± 22.6 | 894.6 ± 65.3 | 0.67 | 0.42 | 0.04 | 0.84 | 0.02 | 0.89 |

Supplementary Table 3

| | | | DEV | CTR | Females DEV | Females CTR | Males DEV | Males CTR | Effect of treatment F | Effect of treatment p* | Effect of sex F | Effect of sex p | Interaction effect F | Interaction effect p |
|---|---|---|---|---|---|---|---|---|---|---|---|---|---|---|
| Nr of instances | Initiator vs responder | Initiator | **236.5** ± 17.8 | **218.6** ± 19.5 | 202.3 ± 16.1 | 207.4 ± 17.0 | 277.6 ± 23.4 | 232.9 ± 38.1 | 0.50 | 0.49 | 3.20 | 0.09 | 0.79 | 0.39 |
| | | Responder | **15.3** ± 3.6 | **15.0** ± 2.5 | 9.8 ± 1.4 | 12.3 ± 2.0 | 21.8 ± 6.6 | 18.4 ± 4.7 | 0.01 | 0.92 | 4.71 | 0.04 | 0.50 | 0.49 |
| | | Ratio initiator | **0.9** ± 0.0 | **0.9** ± 0.0 | 0.9 ± 0.0 | 0.9 ± 0.0 | 0.9 ± 0.0 | 0.9 ± 0.0 | 0.49 | 0.49 | 2.69 | 0.12 | 0.00 | 0.99 |
| | Initiator | Fighting/boxing/wrestling, | **1.3** ± 0.6 | **0.9** ± 0.5 | 1.5 ± 1.0 | 1.3 ± 0.8 | 1.0 ± 0.4 | 0.3 ± 0.2 | 0.29 | 0.60 | 0.90 | 0.35 | 0.11 | 0.74 |
| | | Grooming others | **0.1** ± 0.1 | **0.1** ± 0.1 | 0.0 ± 0.0 | 0.0 ± 0.0 | 0.2 ± 0.2 | 0.1 ± 0.1 | 0.07 | 0.79 | 2.63 | 0.12 | 0.07 | 0.79 |
| | | Sniffing anogenitally | **71.7** ± 11.7 | **64.6** ± 8.1 | 47.7 ± 5.3 | 54.9 ± 3.4 | 100.6 ± 17.8 | 77.1 ± 17.0 | 0.41 | 0.53 | 8.76 | 0.01 | 1.46 | 0.24 |
| | | Sniffing body | **43.5** ± 4.8 | **39.9** ± 5.4 | 31.0 ± 4.1 | 33.7 ± 4.2 | 58.4 ± 2.4 | 48.0 ± 10.4 | 0.30 | 0.59 | 8.61 | 0.01 | 0.84 | 0.37 |
| | | Sniffing nose-to-nose | **111.6** ± 7.9 | **108.5** ± 8.8 | 112.5 ± 10.6 | 111.9 ± 11.9 | 110.6 ± 11.8 | 104.1 ± 12.7 | 0.07 | 0.80 | 0.13 | 0.72 | 0.05 | 0.83 |
| | | All sniffing | **226.8** ± 18.7 | **213.1** ± 19.7 | 191.2 ± 17.6 | 200.4 ± 17.6 | 269.6 ± 24.0 | 229.3 ± 38.1 | 0.29 | 0.59 | 3.50 | 0.07 | 0.75 | 0.40 |
| | | Stand-off/Nose-off | **8.4** ± 1.8 | **4.6** ± 1.2 | 9.7 ± 3.0 | 5.7 ± 1.9 | 6.8 ± 1.6 | 3.1 ± 0.9 | 2.99 | 0.10 | 1.48 | 0.24 | 0.01 | 0.94 |
| | | Initiator Total | **236.5** ± 17.8 | **218.6** ± 19.5 | 202.3 ± 16.1 | 207.4 ± 17.0 | 277.6 ± 23.4 | 232.9 ± 38.1 | 0.50 | 0.49 | 3.20 | 0.09 | 0.79 | 0.39 |
| | Responder | Social investigation | **8.3** ± 2.1 | **9.4** ± 1.3 | 5.7 ± 0.9 | 8.7 ± 1.3 | 11.4 ± 4.0 | 10.3 ± 2.4 | 0.15 | 0.70 | 2.34 | 0.14 | 0.73 | 0.40 |
| | | Fighting/boxing/wrestling, | **1.0** ± 0.5 | **0.8** ± 0.2 | 0.5 ± 0.3 | 0.7 ± 0.3 | 1.6 ± 1.0 | 0.9 ± 0.3 | 0.30 | 0.59 | 1.49 | 0.24 | 0.74 | 0.40 |
| | | Sniffing anogenitally | **0.1** ± 0.1 | **0.3** ± 0.2 | 0.0 ± 0.0 | 0.2 ± 0.1 | 0.2 ± 0.2 | 0.4 ± 0.4 | 0.75 | 0.40 | 0.61 | 0.44 | 0.00 | 0.99 |
| | | Sniffing body | **1.1** ± 0.4 | **0.8** ± 0.3 | 0.7 ± 0.2 | 0.1 ± 0.1 | 1.6 ± 0.9 | 1.6 ± 0.4 | 0.41 | 0.53 | 6.82 | 0.02 | 0.33 | 0.57 |
| | | Sniffing nose-to-nose | **7.1** ± 1.7 | **8.3** ± 1.2 | 5.0 ± 1.0 | 8.3 ± 1.3 | 9.6 ± 3.3 | 8.3 ± 2.1 | 0.23 | 0.64 | 1.17 | 0.29 | 1.22 | 0.28 |
| | | Stand-off/Nose-off | **6.0** ± 1.6 | **4.9** ± 1.4 | 3.7 ± 0.9 | 3.0 ± 1.0 | 8.8 ± 2.8 | 7.3 ± 2.6 | 0.27 | 0.61 | 5.08 | 0.03 | 0.04 | 0.84 |
| | | Resp total | **15.3** ± 3.6 | **15.0** ± 2.5 | 9.8 ± 1.4 | 12.3 ± 2.0 | 21.8 ± 6.6 | 18.4 ± 4.7 | 0.01 | 0.92 | 4.71 | 0.04 | 0.50 | 0.49 |
| | Tactile contact | No of instances with tactile contact | **301.5** ± 22.9 | **286.4** ± 21.4 | 277.3 ± 32.3 | 272.9 ± 20.5 | 330.4 ± 26.9 | 303.9 ± 40.3 | 0.21 | 0.65 | 1.55 | 0.23 | 0.11 | 0.75 |
| | | Not in contact(OA&OA_near_walls) | **363.1** ± 22.9 | **328.2** ± 25.1 | 343.5 ± 34.4 | 297.0 ± 36.7 | 386.6 ± 25.0 | 368.3 ± 25.5 | 0.78 | 0.39 | 2.43 | 0.13 | 0.15 | 0.70 |
| | | Beh-wo-contact(OA&OA_near_walls) | **43.7** ± 5.1 | **40.4** ± 5.4 | 44.7 ± 8.7 | 33.9 ± 6.0 | 42.6 ± 4.0 | 48.9 ± 8.7 | 0.08 | 0.78 | 0.63 | 0.43 | 1.10 | 0.30 |
| | | In OA not engaged in social behavior | **406.8** ± 25.8 | **368.6** ± 29.6 | 388.2 ± 40.7 | 330.9 ± 42.1 | 429.2 ± 25.6 | 417.1 ± 32.1 | 0.66 | 0.43 | 2.22 | 0.15 | 0.28 | 0.60 |

Supplementary Table 3

| | | | | Females | | Males | | Effect of treatment | | Effect of sex | | Interaction effect | |
|---|---|---|---|---|---|---|---|---|---|---|---|---|---|
| | | **DEV** | **CTR** | DEV | CTR | DEV | CTR | F | p* | F | p | F | p |
| Responding by treatment | Initiator tow D | **85.7** ± 13.7 | **104.3** ± 9.0 | 66.5 ± 9.9 | 93.8 ± 7.0 | 108.8 ± 23.8 | 117.7 ± 17.0 | 1.35 | 0.26 | 4.51 | 0.05 | 0.35 | 0.56 |
| | Initiator tow C | **150.7** ± 14.2 | **114.3** ± 14.3 | 135.8 ± 21.2 | 113.7 ± 16.6 | 168.6 ± 14.6 | 115.0 ± 24.9 | 2.87 | 0.10 | 0.58 | 0.45 | 0.49 | 0.49 |
| | Initiator Total | **236.5** ± 17.8 | **218.5** ± 19.5 | 202.3 ± 16.1 | 207.4 ± 17.0 | 277.4 ± 23.3 | 232.7 ± 38.1 | 0.49 | 0.49 | 3.18 | 0.09 | 0.78 | 0.39 |
| | Responder tow D | **5.0** ± 1.0 | **7.8** ± 1.4 | 3.3 ± 1.2 | 5.7 ± 0.9 | 7.0 ± 0.8 | 10.4 ± 2.7 | 2.53 | 0.13 | 5.41 | 0.03 | 0.09 | 0.77 |
| | Responder tow C | **10.3** ± 3.1 | **7.2** ± 1.2 | 6.5 ± 1.5 | 6.7 ± 1.4 | 14.8 ± 6.1 | 7.9 ± 2.1 | 1.25 | 0.27 | 2.46 | 0.13 | 1.38 | 0.25 |
| | Responder Total | **15.3** ± 3.6 | **14.9** ± 2.4 | 9.8 ± 1.4 | 12.3 ± 2.0 | 21.8 ± 6.6 | 18.3 ± 4.6 | 0.02 | 0.90 | 4.74 | 0.04 | 0.53 | 0.47 |
| | Adjusted Initiator tow D | **39.9** ± 4.7 | **38.6** ± 5.0 | 31.6 ± 3.4 | 33.8 ± 3.1 | 49.9 ± 7.4 | 44.9 ± 10.1 | 0.04 | 0.85 | 4.23 | 0.05 | 0.25 | 0.63 |
| | Adjusted Initiator tow C | **39.1** ± 2.5 | **34.8** ± 2.7 | 34.3 ± 2.9 | 35.3 ± 2.7 | 44.9 ± 2.6 | 34.3 ± 5.0 | 1.50 | 0.23 | 1.48 | 0.24 | 2.17 | 0.15 |
| | Adjusted Responder tow D | **2.6** ± 0.6 | **2.9** ± 0.5 | 1.6 ± 0.5 | 2.2 ± 0.4 | 3.8 ± 0.9 | 3.8 ± 0.9 | 0.16 | 0.70 | 6.60 | 0.02 | 0.16 | 0.69 |
| | Adjusted Responder tow C | **2.6** ± 0.8 | **2.3** ± 0.4 | 1.6 ± 0.3 | 2.1 ± 0.3 | 3.7 ± 1.5 | 2.6 ± 0.7 | 0.18 | 0.68 | 2.78 | 0.11 | 1.06 | 0.32 |
| Nr of instances | Social investigation | **96.7** ± 10.9 | **104.2** ± 10.2 | 109.0 ± 15.5 | 125.9 ± 12.2 | 82.0 ± 20.8 | 76.3 ± 10.0 | 0.16 | 0.70 | 7.38 | 0.01 | 0.64 | 0.43 |
| | Non-social invest | **156.9** ± 19.2 | **161.1** ± 13.9 | 199.5 ± 27.8 | 199.8 ± 12.2 | 105.8 ± 25.7 | 111.4 ± 11.6 | 0.03 | 0.86 | 31.25 | 0.00 | 0.03 | 0.87 |
| | Interacting with the | **2.9** ± 0.9 | **3.7** ± 1.1 | 3.3 ± 0.9 | 4.0 ± 1.7 | 2.4 ± 1.7 | 3.3 ± 1.1 | 0.22 | 0.64 | 0.25 | 0.62 | 0.00 | 0.95 |
| | Fighting/boxing/wrestling /kicking | **0.8** ± 0.4 | **0.5** ± 0.2 | 1.0 ± 0.6 | 0.7 ± 0.3 | 0.6 ± 0.4 | 0.3 ± 0.2 | 0.58 | 0.45 | 0.85 | 0.37 | 0.00 | 0.98 |
| | Following | **3.4** ± 1.0 | **2.8** ± 0.9 | 2.3 ± 0.7 | 1.7 ± 0.6 | 4.6 ± 2.0 | 4.3 ± 1.7 | 0.13 | 0.72 | 3.16 | 0.09 | 0.02 | 0.90 |
| | Any other behavior | **0.3** ± 0.2 | **0.1** ± 0.1 | 0.0 ± 0.0 | 0.0 ± 0.0 | 0.6 ± 0.4 | 0.3 ± 0.2 | 0.79 | 0.38 | 6.28 | 0.02 | 0.79 | 0.38 |
| Burrow | Non-social exploration | **17.3** ± 4.0 | **16.9** ± 2.2 | 27.0 ± 4.0 | 22.3 ± 2.1 | 5.6 ± 1.8 | 9.9 ± 2.2 | 0.01 | 0.95 | 33.59 | 0.00 | 2.33 | 0.14 |
| | Passive/ immobile alone | **6.5** ± 1.2 | **7.9** ± 1.4 | 8.8 ± 0.9 | 9.9 ± 2.0 | 3.8 ± 1.8 | 5.3 ± 1.3 | 0.45 | 0.51 | 6.49 | 0.02 | 0.01 | 0.91 |
| | Passive/immobile social | **0.6** ± 0.2 | **1.9** ± 0.4 | 0.8 ± 0.3 | 2.6 ± 0.6 | 0.4 ± 0.4 | 1.0 ± 0.4 | 4.321 | 0.049 (0.15) | 3.17 | 0.09 | 1.01 | 0.33 |
| | Rearing supported | **4.4** ± 1.3 | **3.2** ± 1.2 | 7.2 ± 1.7 | 5.7 ± 1.7 | 1.0 ± 0.5 | 0.0 ± 0.0 | 0.66 | 0.42 | 14.88 | 0.00 | 0.03 | 0.87 |
| | Rearing unsupported | **0.3** ± 0.3 | **0.3** ± 0.1 | 0.5 ± 0.5 | 0.4 ± 0.2 | 0.0 ± 0.0 | 0.0 ± 0.0 | 0.01 | 0.92 | 2.82 | 0.11 | 0.01 | 0.92 |
| | Self grooming | **10.8** ± 2.8 | **14.7** ± 2.7 | 15.5 ± 4.0 | 19.1 ± 3.2 | 5.2 ± 1.8 | 9.0 ± 3.5 | 0.97 | 0.34 | 7.33 | 0.01 | 0.00 | 0.98 |
| | Sniffing anogenitally | **27.8** ± 4.8 | **29.9** ± 3.0 | 25.3 ± 2.8 | 34.7 ± 3.1 | 30.8 ± 9.9 | 23.7 ± 4.7 | 0.04 | 0.84 | 0.24 | 0.63 | 2.16 | 0.16 |
| | Sniffing body | **9.6** ± 1.6 | **9.8** ± 1.5 | 12.0 ± 2.1 | 13.0 ± 2.0 | 6.8 ± 1.9 | 5.6 ± 0.9 | 0.00 | 0.96 | 9.88 | 0.01 | 0.31 | 0.59 |
| | Sniffing nose-to-nose | **59.3** ± 6.7 | **64.6** ± 6.6 | 71.7 ± 5.6 | 78.2 ± 8.3 | 44.4 ± 9.7 | 47.0 ± 5.9 | 0.29 | 0.60 | 11.76 | 0.00 | 0.05 | 0.82 |
| | All sniffing | **96.7** ± 10.9 | **104.2** ± 10.2 | 109.0 ± 6.7 | 125.9 ± 12.2 | 82.0 ± 20.8 | 76.3 ± 10.0 | 0.16 | 0.70 | 7.38 | 0.01 | 0.64 | 0.43 |
| | Stand-off/Nose-off | **8.7** ± 2.2 | **6.3** ± 1.5 | 10.0 ± 3.7 | 6.3 ± 2.1 | 7.2 ± 1.7 | 6.3 ± 2.2 | 0.68 | 0.42 | 0.26 | 0.61 | 0.25 | 0.63 |
| | Walking/running | **139.6** ± 16.0 | **144.3** ± 12.1 | 172.5 ± 6.9 | 177.4 ± 11.2 | 100.2 ± 24.4 | 101.6 ± 9.6 | 0.05 | 0.42 | 25.93 | 0.00 | 0.02 | 0.90 |
| | Burrow Total | **292.4** ± 33.4 | **306.6** ± 25.7 | 358.0 ± 18.8 | 376.0 ± 24.8 | 213.6 ± 51.0 | 217.4 ± 20.3 | 0.12 | 0.42 | 23.00 | 0.00 | 0.05 | 0.82 |

Supplementary Table 3

| | | | | | Females | | Males | | Effect of treatment | | Effect of sex | | Interaction effect | |
|---|---|---|---|---|---|---|---|---|---|---|---|---|---|---|
| | | | **DEV** | **CTR** | DEV | CTR | DEV | CTR | F | p* | F | p | F | p |
| **Nr of instances** | **Full open area** | Social investigation | **133.5** ± 20.3 | **118.6** ± 18.4 | 80.3 ± 13.0 | 81.9 ± 13.9 | 197.4 ± 17.9 | 165.9 ± 29.8 | 0.47 | 0.50 | 20.93 | 0.00 | 0.57 | 0.46 |
| | | Non-social invest | **355.8** ± 24.7 | **319.1** ± 25.4 | 349.7 ± 54.6 | 295.4 ± 35.3 | 363.2 ± 21.8 | 349.6 ± 33.0 | 0.75 | 0.40 | 0.75 | 0.40 | 0.27 | 0.61 |
| | | Interacting with the envirc | **3.4** ± 1.1 | **3.0** ± 0.7 | 4.7 ± 1.7 | 3.6 ± 1.2 | 1.8 ± 0.6 | 2.3 ± 0.6 | 0.06 | 0.82 | 2.46 | 0.13 | 0.37 | 0.55 |
| | | Fighting/boxing/wrestling, | **1.5** ± 0.7 | **1.4** ± 0.6 | 1.0 ± 0.7 | 1.4 ± 0.8 | 2.0 ± 1.2 | 1.4 ± 0.9 | 0.00 | 0.95 | 0.25 | 0.63 | 0.26 | 0.61 |
| | | Following | **11.9** ± 4.6 | **11.1** ± 3.6 | 3.5 ± 1.5 | 2.0 ± 0.6 | 22.0 ± 7.8 | 22.7 ± 5.6 | 0.01 | 0.93 | 18.06 | 0.00 | 0.06 | 0.81 |
| | | Grooming others | **0.2** ± 0.1 | **0.1** ± 0.1 | 0.0 ± 0.0 | 0.0 ± 0.0 | 0.4 ± 0.2 | 0.1 ± 0.1 | 1.19 | 0.29 | 5.31 | 0.03 | 1.19 | 0.29 |
| | | Hiding alone | **7.5** ± 1.9 | **8.4** ± 2.7 | 11.7 ± 2.2 | 12.8 ± 4.1 | 2.6 ± 1.2 | 2.7 ± 0.9 | 0.03 | 0.86 | 8.37 | 0.01 | 0.02 | 0.88 |
| | | Hiding social | **10.4** ± 2.6 | **6.0** ± 1.2 | 15.0 ± 3.7 | 7.8 ± 1.8 | 4.8 ± 1.2 | 3.7 ± 1.1 | 3.05 | 0.09 | 8.98 | 0.01 | 1.66 | 0.21 |
| | | Any other behavior | **6.4** ± 3.2 | **3.1** ± 1.1 | 0.2 ± 0.2 | 0.1 ± 0.1 | 13.8 ± 5.4 | 7.0 ± 1.6 | 2.02 | 0.17 | 18.14 | 0.00 | 1.96 | 0.18 |
| | | Non-social exploration | **26.0** ± 2.4 | **27.0** ± 3.9 | 26.0 ± 4.1 | 22.3 ± 3.6 | 26.0 ± 2.2 | 33.0 ± 6.9 | 0.10 | 0.76 | 1.02 | 0.32 | 1.02 | 0.32 |
| | | Passive/ immobile alone | **3.6** ± 1.4 | **2.1** ± 0.7 | 1.0 ± 0.4 | 0.4 ± 0.2 | 6.8 ± 2.3 | 4.3 ± 1.2 | 1.69 | 0.21 | 16.69 | 0.00 | 0.69 | 0.42 |
| | | Passive/immobile social | **0.3** ± 0.1 | **0.6** ± 0.2 | 0.3 ± 0.2 | 0.4 ± 0.3 | 0.2 ± 0.2 | 0.9 ± 0.4 | 1.27 | 0.27 | 0.17 | 0.69 | 0.64 | 0.43 |
| | | Rearing supported | **50.8** ± 4.5 | **49.2** ± 4.5 | 60.2 ± 4.6 | 50.1 ± 7.3 | 39.6 ± 4.5 | 48.0 ± 4.3 | 0.02 | 0.90 | 2.93 | 0.10 | 1.94 | 0.18 |
| | | Rearing unsupported | **10.7** ± 2.6 | **8.3** ± 2.5 | 13.0 ± 3.8 | 7.6 ± 2.1 | 8.0 ± 2.8 | 9.3 ± 5.0 | 0.28 | 0.60 | 0.17 | 0.68 | 0.72 | 0.40 |
| | | Self grooming | **24.9** ± 6.3 | **22.9** ± 4.0 | 11.0 ± 3.0 | 10.8 ± 2.5 | 41.6 ± 8.8 | 38.6 ± 3.3 | 0.12 | 0.73 | 38.24 | 0.00 | 0.09 | 0.77 |
| | | Sniffing anogenitally | **44.5** ± 8.7 | **36.6** ± 7.2 | 22.3 ± 3.5 | 21.0 ± 3.5 | 71.2 ± 9.4 | 56.7 ± 12.3 | 0.84 | 0.37 | 23.96 | 0.00 | 0.58 | 0.45 |
| | | Sniffing body | **34.3** ± 5.9 | **30.9** ± 5.6 | 18.3 ± 4.3 | 20.6 ± 3.3 | 53.4 ± 2.9 | 44.3 ± 9.9 | 0.27 | 0.61 | 19.55 | 0.00 | 0.73 | 0.40 |
| | | Sniffing nose-to-nose | **54.5** ± 9.3 | **51.0** ± 6.5 | 39.7 ± 4.5 | 40.3 ± 8.0 | 72.4 ± 16.6 | 64.7 ± 8.3 | 0.11 | 0.74 | 7.46 | 0.01 | 0.16 | 0.69 |
| | | All sniffing | **133.4** ± 20.2 | **118.6** ± 18.4 | 80.3 ± 11.3 | 81.9 ± 13.9 | 197.0 ± 17.7 | 165.7 ± 29.7 | 0.46 | 0.51 | 20.88 | 0.00 | 0.56 | 0.46 |
| | | Stand-off/Nose-off | **6.5** ± 1.6 | **4.8** ± 1.2 | 3.8 ± 0.6 | 3.1 ± 1.0 | 9.8 ± 2.9 | 7.0 ± 2.2 | 0.87 | 0.36 | 6.79 | 0.02 | 0.30 | 0.59 |
| | | Walking/running | **329.8** ± 23.0 | **292.1** ± 22.3 | 323.7 ± 37.7 | 273.1 ± 32.2 | 337.2 ± 22.4 | 316.6 ± 27.1 | 1.02 | 0.32 | 0.66 | 0.43 | 0.18 | 0.68 |
| | | Full OA Total | **627.4** ± 48.4 | **558.8** ± 50.8 | 555.5 ± 65.4 | 477.4 ± 61.8 | 713.6 ± 49.2 | 663.3 ± 66.1 | 0.83 | 0.37 | 5.94 | 0.02 | 0.04 | 0.85 |

*in parenthesis, p corrected with B-H procedure where applicable

C - CTR - Control

D - DEV - Devocalized

Supplementary Table 4

| | | | DEV | CTR | Females DEV | Females CTR | Males DEV | Males CTR | Effect of treatment F | Effect of treatment p | Effect of sex F | Effect of sex p | Interaction effect F | Interaction effect p |
|---|---|---|---|---|---|---|---|---|---|---|---|---|---|---|
| Latencies | Whole environment | Non-social exploration | **63.5** ± 15.4 | **86.6** ± 26.9 | 37.1 ± 12.5 | 64.4 ± 17.1 | 95.2 ± 23.5 | 115.2 ± 55.7 | 0.43 | 0.52 | 2.25 | 0.15 | 0.01 | 0.92 |
| | | Rearing supported | **85.1** ± 25.1 | **125.2** ± 29.3 | 63.5 ± 23.0 | 127.3 ± 43.6 | 111.1 ± 45.2 | 122.5 ± 36.8 | 0.72 | 0.40 | 0.23 | 0.63 | 0.35 | 0.56 |
| | | Rearing unsupported | **1427.2** ± 304.8 | **1271.0** ± 313.4 | 1328.7 ± 525.2 | 1120.7 ± 457.0 | 1545.3 ± 218.0 | 1464.3 ± 397.7 | 0.09 | 0.77 | 0.33 | 0.57 | 0.02 | 0.90 |
| | | Self grooming | **260.9** ± 64.5 | **203.9** ± 51.0 | 349.7 ± 88.5 | 211.2 ± 79.6 | 154.3 ± 68.4 | 194.5 ± 55.6 | 0.34 | 0.57 | 1.57 | 0.22 | 1.11 | 0.30 |
| | | Any rearing | **84.2** ± 25.3 | **122.1** ± 29.8 | 61.8 ± 23.6 | 121.7 ± 44.5 | 111.1 ± 45.2 | 122.5 ± 36.8 | 0.64 | 0.43 | 0.31 | 0.58 | 0.29 | 0.59 |
| | | Any Sniffing | **11.1** ± 3.5 | **10.2** ± 3.3 | 12.8 ± 5.2 | 13.4 ± 5.3 | 9.2 ± 4.2 | 6.2 ± 2.4 | 0.05 | 0.82 | 1.07 | 0.31 | 0.12 | 0.74 |

The role of rat ultrasonic vocalizations on social and non-social investigation in the seminatural environment

**Supplementary Table 5. Benjamini-Hochberg correction**

| | Behavior upon rank p-value from large to small | Original p-value | Rank | Nr of comparisons | Corrected p-value | Final p-value |
|---|---|---|---|---|---|---|
| Duration | Nose-off (OA Centre) | **0.020** | 1 | 8 | 0.16 | 0.10 |
| | Passive immobile alone (In opening) | **0.030** | 2 | 8 | 0.12 | 0.10 |
| | Fighting/boxing/wrestling/kicking towards CTR | **0.036** | 3 | 8 | 0.10 | 0.10 |
| | Any other behavior | **0.050** | 4 | 8 | 0.10 | 0.10 |
| | Self-grooming | 0.350 | 5 | 8 | 0.56 | 0.56 |
| | Conflict behavior | 0.530 | 6 | 8 | 0.71 | 0.65 |
| | Non-social exploration | 0.570 | 7 | 8 | 0.65 | 0.65 |
| | All sniffing | 0.840 | 8 | 8 | 0.84 | 0.84 |
| Nr of episodes | passive immobile soc | **0.020** | 1 | 6 | 0.12 | 0.12 |
| | passive immobile soc (Burrow) | **0.049** | 2 | 6 | 0.15 | 0.15 |
| | conflict behavior | 0.291 | 3 | 6 | 0.58 | 0.58 |
| | all sniffing | 0.679 | 4 | 6 | 1.02 | 0.78 |
| | self-grooming | 0.689 | 5 | 6 | 0.83 | 0.78 |
| | non-social exploration | 0.782 | 6 | 6 | 0.78 | 0.78 |

We used predetermined behaviors: all sniffing, non-social exploration, self-grooming and conflict behavior.
According to the Benjamini-Hochberg procedure:
For corrected p-value we multiplied the original p-value by the quotient of Nr of comparisons/rank
Final p-value is the smaller of the two: either corrected p-value or final p-value of the next rank
